# WEE1 kinase inhibition triggers severe chromosome pulverization in aneuploid cells

**DOI:** 10.1101/2023.09.19.558475

**Authors:** Maria M. Haykal, Sylvie Rodrigues-Ferreira, Clara Nahmias

## Abstract

Aneuploidy, a hallmark of cancer, is a prominent feature associated with poor prognosis in breast cancer. Here, we screened a panel of cell cycle kinase inhibitors to identify novel targets for highly aneuploid breast cancers. We show that increasing aneuploidy in breast cancer cells sensitizes to the inhibition of WEE1 kinase. Upon exposure to WEE1 inhibitor, aneuploid cells exhibit aberrant mitosis characterized by the detachment of centromere proteins from centromeric DNA and pulverization of chromosomes. The occurrence of such phenotype is driven by excessive levels of replication stress and DNA damage during S-phase, that in turn trigger major defects in the subsequent mitosis. We show that DNA2 helicase/nuclease, that regulates replication of centromeric DNA, is the key player responsible for severe chromosome pulverization in mitosis. The heightened vulnerability of aneuploid cells to WEE1 inhibition, coupled with underlying molecular mechanisms, provides a rationale for clinical exploration of WEE1-targeted therapies against aneuploid breast cancers.

**Impact Statement:** Increased vulnerability of aneuploid cells to WEE1 inhibition is orchestrated by the DNA2 nuclease/helicase. These findings open new therapeutic strategies in the context of personalized medicine in breast cancer.

## Introduction

Cellular DNA content is carefully controlled by cell cycle checkpoints, which monitor and regulate DNA replication and subsequent cell division, preventing genomic abnormalities^1^. Chromosomal instability (CIN), a condition of persistent chromosome missegregation, is a significant hallmark of human cancers^2^. Changes in mitotic spindle assembly checkpoint function, centrosome duplication, kinetochore function, and microtubule stability have all been linked to CIN^3^. Aneuploidy, an inevitable consequence of CIN, refers to the condition in which cells contain an atypical quantity of chromosomes. Aneuploidy is detected in 90% of solid tumors^4^. The occurrence of aneuploidy generally reduces a cell’s proliferative ability owing to the detrimental effects of proteotoxic stress and induction of DNA damage^5–7^. However, in tumors, high levels of aneuploidy are strongly associated with poor patient outcomes, suggesting that they may confer a growth advantage and contribute to some aspects of cancer^8^.

Breast cancer is a heterogeneous disease affecting women worldwide. It is the most frequently diagnosed cancer and a leading cause of cancer-related deaths^9^. Breast cancer genomes are often aneuploid and display intricate numerical and structural chromosomal rearrangements^10–12^. Importantly, high levels of aneuploidy are associated with a poor prognosis in the majority of breast cancer cases^13–15^. Given the unfavorable prognosis associated with aneuploidy levels in breast cancer, there is an urgent need to design adapted therapies for these tumors^16,17^.

ATIP3, a microtubule-stabilizing protein, is a prognostic biomarker for breast cancer patient survival^18,19^. Recent work has shown that depletion of ATIP3 induces centrosome amplification and the formation of multipolar mitotic spindles, thereby increasing aneuploidy. Interestingly, ATIP3 deficiency in breast tumors also predicts their response to taxanes: tumors expressing low levels of ATIP3 are associated with high CIN and are aneuploid, but paradoxically more responsive to taxane-based chemotherapy^17^. This phenomenon can be attributed to the detrimental levels of excessive aneuploidy resulting from combined ATIP3 depletion and taxane treatment, ultimately driving cell death^20,21^. Nonetheless, taxanes are highly toxic drugs with adverse side effects, highlighting the need for targeted therapies.

In the search for new targeted therapies for highly aneuploid breast cancers, we depleted ATIP3 as a model to increase aneuploidy levels, and screened a panel of kinase inhibitors known to perturb the cell cycle. In this study, we focused on WEE1 kinase, a gatekeeper of the G2/M cell cycle checkpoint^22^. By inhibiting cyclin-dependent kinases CDK1 and CDK2 through phosphorylation of Tyr15 during G2/M and S phase, respectively^23^. WEE1 safeguards DNA replication^24–26^ and prevents cells from entering mitosis until DNA replication and repair processes are completed, thus maintaining genome stability. However, it remains unclear whether WEE1 effects occurring in S phase and in mitosis may be directly related. Here we show that in highly aneuploid cancer cells, that already harbor DNA alterations^27^, inhibiting WEE1 kinase elevates the levels of replication stress and DNA damage in S-phase, which in turn are responsible for severe chromosome pulverization by DNA2 nuclease in subsequent mitosis, causing massive cell death.

## Results

The search for new therapeutic strategies against highly aneuploid breast cancers led us to perform a chemical synthetic lethality screen of 28 cell cycle kinase inhibitors in breast cancer cells in which ATIP3 protein was depleted to increase aneuploidy. The SUM52PE breast cancer cell line expressing or not ATIP3 was grown in 3-dimensions as multicellular spheroids (MCSs) to mimic the features of solid tumors and was treated with increasing doses of each inhibitor (Figure 1A). After 72 h of treatment, cell viability was assessed and the IC50 of each inhibitor was calculated. We considered an inhibitor as a differential hit if the IC50-fold change between control and ATIP3-depleted cells was equal to or higher than 2 (Figure 1B, Figure 1-supplement Table S1). ATIP3 depletion improved the cytotoxic response to six different inhibitors targeting ATR, ATM, WEE1, Aurora, and PLK4 kinases (Figure 1-figure supplement 1A,B,C). Among these differential inhibitors, we focused on AZD1775, which is a WEE1 kinase inhibitor. AZD1775 was more potent in ATIP3-depleted breast cancer MCSs, as indicated by their size (Figure 1C) and lower IC50 values (mean IC50 in shCtl is 1.31 µM *vs*. 0.43 µM in shATIP3) (Figure 1D). AZD1775 also exhibited lower IC50 values in two other breast cancer MCSs models in which ATIP3 expression was depleted (Figure 1-figure supplement 1D). Similar results were obtained with PD0166285, another WEE1 inhibitor (Figure 1-figure supplement 1E). In addition, AZD1775 abolished the phosphorylation of CDK1 at Tyr15, confirming target engagement (Figure 1-figure supplement1F). Importantly, the doses of AZD1775 that caused maximal cell death in tumor cells had very little effects on diploid, non-transformed RPE-1 cells (Figure 1E). We then investigated whether increased vulnerability of ATIP3-depleted aneuploid cells to WEE1 inhibition was driven by aneuploidy *per se* or by ATIP3-specific effects. To address this question, we induced aneuploidy in RPE-1 cells using reversine, an inhibitor of MPS1 kinase that induces chromosome missegregation^28,29^, and combined it with WEE1 inhibition (Figure 1-figure supplement 1G). MPS1 inhibition in RPE-1 cells resulted in aneuploidy, as shown by increased variability in chromosome numbers (Figure 1-figure supplement 1H). Aneuploidy induction by reversine in RPE-1 cells rendered them vulnerable to WEE1 inhibition, leading to cell death (Figure 1F), indicating that sensitivity to WEE1 inhibition is associated with aneuploidy. We then tested the efficacy of AZD1775 *in vivo* using xenografts of the MDA-MB-468 breast cancer cell line in which ATIP3 was depleted. AZD1775 was administered daily by oral gavage at a dose of 90 mg/kg for 3 weeks. ATIP3 depletion increased tumor growth, in line with the high aggressiveness of tumors expressing low levels of ATIP3 (Figure 1G). Treatment with AZD1775 prevented tumor growth and had a more prominent effect on ATIP3-depleted tumors (Fold change in tumor volume of 1.6 in shCtl *vs*. shCtl AZD1775 and 2.3 in shATIP3 *vs*. shATIP3 AZD1775) (Figure 1H) in agreement with *in vitro* observations.

**Figure 1:**
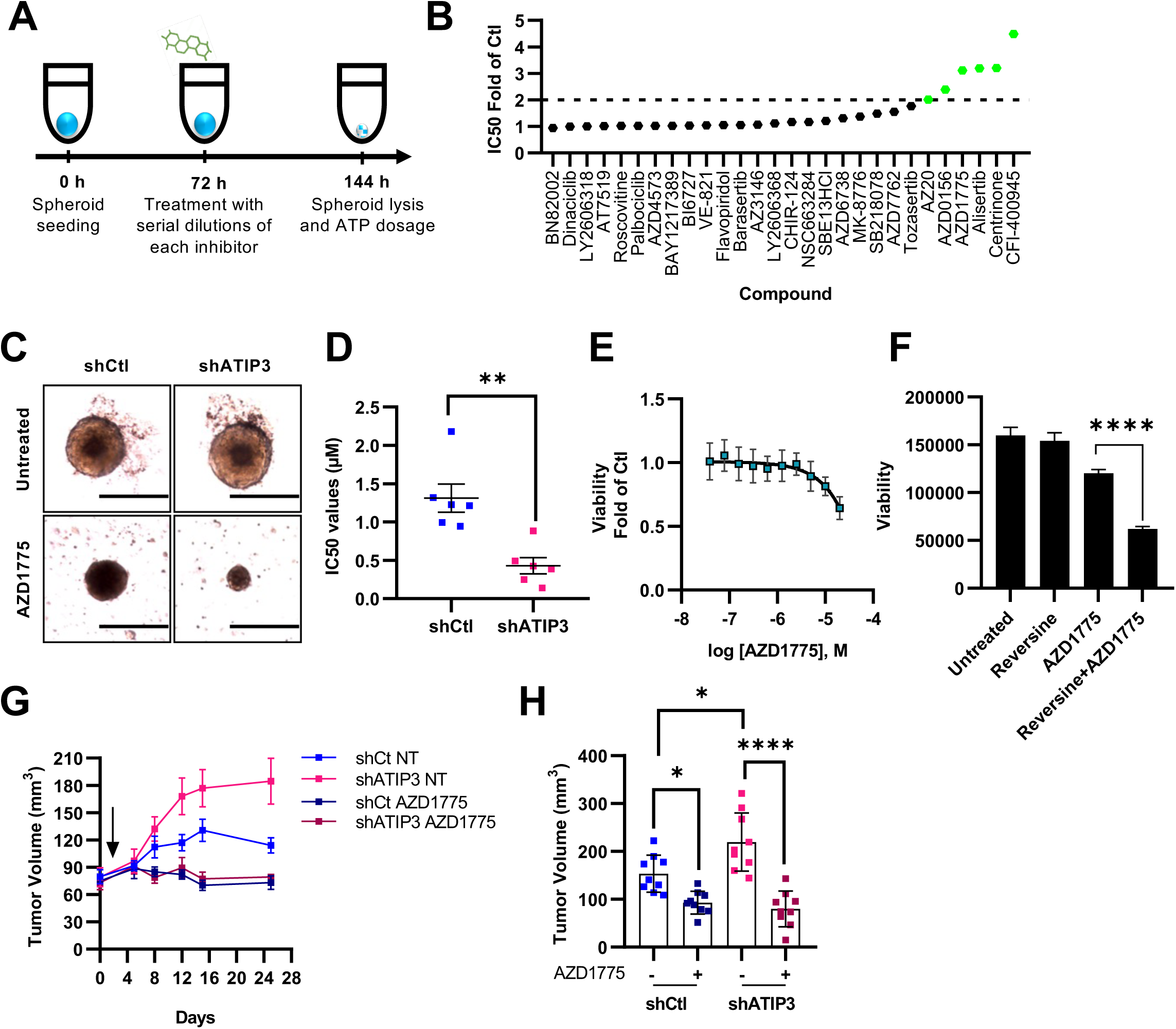
Chemical synthetic screen identifies WEE1 kinase as a target for ATIP3-deficient breast cancer cells. **(A)** Pipeline scheme for drug screening using 28 cell cycle kinase inhibitors. The SUM52PE cell was cultured as MSCs. Screening was conducted in quadruplicates. **(B)** Plot representing the IC50 fold change of each inhibitor between shCtl and shATIP3. **(C)** Representative images of SUM52PE MCSs treated or not with 500 nM AZD1775 for 72 h *(Magnification 20x)*. **(D)** IC50 values of AZD1775 in MCSs (mean ± SEM. of N=6; two-tailed t-test; **p<0.01). **(E)** Plot representing viability of RPE-1 cells upon 72 h of treatment with increasing doses of AZD1775 (mean ± SEM. of N=3). **(F)** Plot representing viability of RPE-1 cells pre-treated with reversine for 48 h and 10 µM AZD1775 for 72 h (mean ± S.D.; one-way ANOVA; ****p<0.0001). **(G-H)** MDA-MB-468 shCtl or shATIP3 subcutaneous xenografts were treated or not with 90 mg/kg of AZD1775 by oral gavage (mean ± SEM; N=1 with 9 mice per group) **(G)** Tumor growth curve (the beginning of treatment is indicated by an arrow). **(H)** Tumor volume (mm^3^) after 15 days of treatment (one-way ANOVA; *p<0.05 ****p<0.0001).

WEE1 inhibition has been shown to induce premature mitotic entry of S-phase-arrested cells after induction of DNA damage^30^. To evaluate mitotic entry, we used live cell microscopy on control or ATIP3-depleted cells (Figure 2A, and Figure 2-figure supplement 1A) that were synchronized at the G1/S boundary and released in the presence or absence of AZD1775. Interestingly, in response to WEE1 inhibition, ATIP3-depleted cells entered mitosis an hour earlier than control cells (3.1 h post release for siATIP3 to enter mitosis vs. 4.2 h for siCtl), whereas a majority of untreated cells did not enter mitosis at 6 h post-release (Figure 2B). In addition, WEE1 inhibition increased the phosphorylation of both histone H3 and CDK1 substrates (Figure 2-figure supplement 1B,C), indicators of mitotic entry, in ATIP3 depleted cells. In line with these observations, the proportion of mitotic cells increased in ATIP3-depleted cells in both 2-dimensions (Figure 2C) and MCSs (Figure 2-figure supplement 1D,E). To gain further insights into the effects induced by WEE1 inhibition in mitosis, we filmed HeLa cells stably expressing mCherry-histone H2B in which we depleted ATIP3 to closely track the fate of individual cells. As described previously^17^, untreated ATIP3-depleted cells spent more time in mitosis compared to control cells (Figure 2D,E, and Movies S1 and S2). WEE1 inhibition further prolonged the time spent in mitosis. Notably, ATIP3-depleted cells treated with AZD1775 experienced significantly longer mitosis than control cells (Figure 2D,E, and Movies S3 and S4). Strikingly, 75% of ATIP3-depleted cells treated with AZD1775 exhibited a back-and-forth movement of their DNA around the mitotic spindle, compared to only 1% of control cells that behaved in this manner (Figure 2-figure supplement 1F, and Movie S4), which is consistent with exceedingly long mitosis. AZD1775 induced two types of cell death. Cells either died during mitosis (mitotic catastrophe) (Figure 2D panel 3, and Movie S3) or after mitotic exit (Figure 2D panel 4, and Movie S4). Of note, cell death occurred in 95% of ATIP3-depleted cells treated with AZD1775 (63% after mitotic exit) and in 78% of control cells (12% after mitotic exit) (Figure 2F,G). Together, these results show that ATIP3 depletion accelerates mitotic entry, prolongs the time spent in mitosis, and exacerbates cell death upon exposure to WEE1 inhibitor.

**Figure 2:**
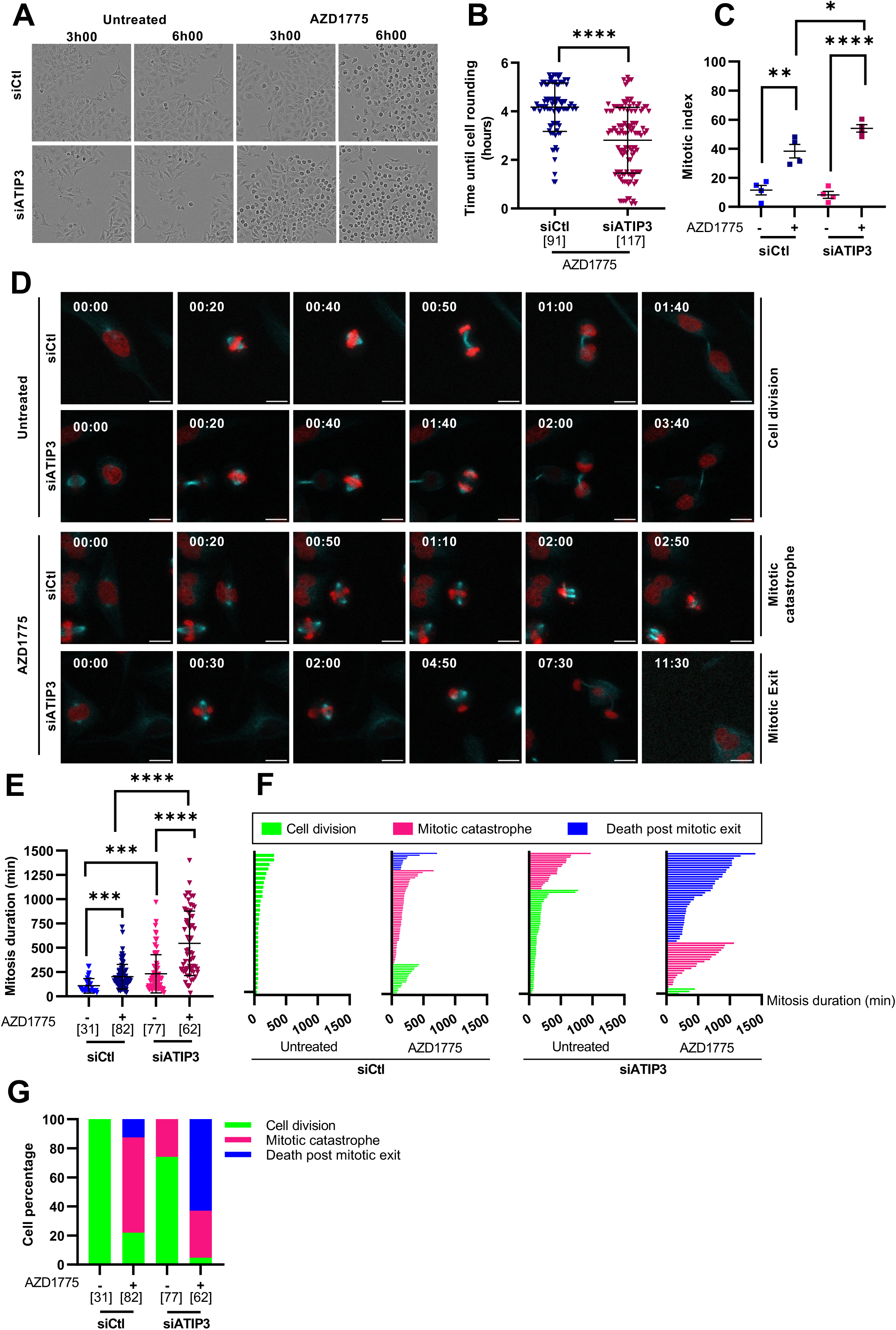
WEE1 inhibition accelerates mitotic entry in aneuploid cells and induces their cell death after failed mitosis. **(A-B)** HeLa cells expressing or not ATIP3 were synchronized at the G1/S boundary, released in the presence or absence of 500 nM AZD1775, and filmed for 6 h. **(A)** Representative images *(Magnification 10x)*. **(B)** Scattered dot plot showing the time to enter mitosis in response to AZD1775 (elapsed time between release and cell rounding). The number of analyzed cells is in brackets (mean ± S.D.; Mann-Whitney test; ****p<0.0001). **(C)** HeLa cells expressing or not ATIP3 were treated or not with 500 nM AZD1775 for 6 h. Mitotic index was determined as the ratio of mitotic cells to the cellular population (mean ± SEM. of N=4; a minimum of 177 cells were analyzed per group; one-way ANOVA; *p<0.05, **p<0.01, ****p<0.0001). **(D-G)** HeLa mCherry-H2B expressing not ATIP3 were stained with siR-tubulin and treated or not with 500 nM AZD1775, then analyzed by timelapse fluorescent imaging for 48 h. **(D)** Representative images. Time is indicated on the top left as h:min. Microtubules (siR-tubulin) are shown in cyan and DNA (mCherry-H2B) in red. **(E)** Scattered dot plot showing the duration of mitosis. The number of analyzed cells is in brackets (mean ± S.D.; Kruskal-Wallis test followed by Dunn’s multiple comparisons; ***p< 0.001, ****p<0.0001). **(F)** Cell fate profiles. **(G)** Proportion of cell fates measured in (F). Scale bar = 20 µm.

WEE1 inhibition caused a very particular phenotype in mitosis, where a bulk of chromatin mass was on the outside of the mitotic spindle rather than in structured chromosomes aligned on the metaphase plate in both 2-dimensions (Figure 3A *enlarged panels*) and MCSs (Figure 2-figure supplement 1G). ATIP3 depletion exacerbated these mitotic abnormalities (Figure 3B). Similar results were obtained when WEE1 was depleted using siRNA, confirming that WEE1 kinase is the main target of AZD1775 (Figure 2-figure supplement 1H). We then investigated the impact of CDK1 and CDK2, the major kinases controlled by WEE1^31^. The abnormal mitosis phenotype was completely abolished upon combining WEE1 inhibition with RO-3306, a CDK1 inhibitor (Figure 2-figure supplement 1I,J) but not following CDK2 depletion (Figure 2-figure supplement 1K), confirming that the abnormal phenotype was mainly due to the activation of CDK1 after WEE1 inhibition.

**Figure 3:**
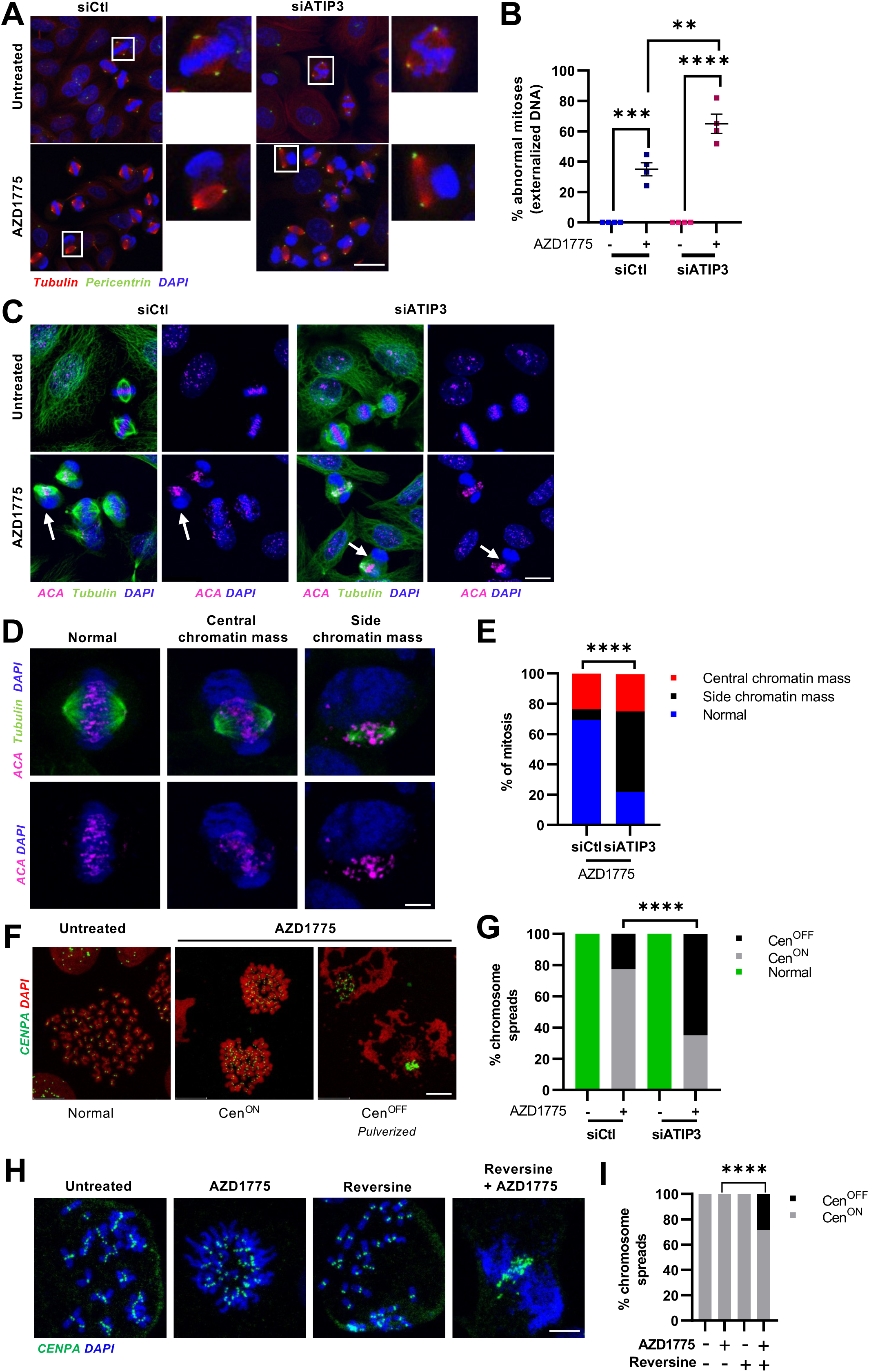
WEE1 inhibition causes severe mitotic defects and chromosome pulverization in highly aneuploid cells. HeLa cells expressing or not ATIP3 were treated or not with 500 nM AZD1775 for 6 h. **(A)** Immunofluorescence representative images showing centrosomes (pericentrin) in green, microtubules (tubulin) in red and DNA (DAPI) in blue. **(B)** Quantification of abnormal mitosis shown in (A) (mean ± SEM. of N=4; a minimum of 128 cells was analyzed; one-way ANOVA; **p<0.01, ***p<0.001, ****p<0.0001). **(C)** Immunofluorescence representative images showing CENPs (ACA) in magenta, microtubules (tubulin) in green and DNA (DAPI) in blue. Scale bar = 20 µm. **(D)** Immunofluorescence representative images of the mitotic phenotypes showing CENPs in magenta, microtubules in green and DNA in blue. Scale bar = 5 µm. **(E)** Quantification of the proportions of the mitotic phenotypes shown in (D) (mean ± SEM. of N=4; a minimum of 112 cells were analyzed per group; two-way ANOVA; side chromatin mass in siCtl *vs*. siATIP3 ****p<0.0001). **(F)** Immunofluorescence representative images of chromosome spreads showing CENP-A in green and DNA in red. **(G)** Quantification of the proportions of chromosome spreads shown in (F) (mean ± SEM. of N=6; a minimum of 145 spreads were analyzed per group; two-way ANOVA; siCtl Cen^OFF^ *vs*. siATIP3 Cen^OFF^ ****p<0.0001). **(H-I)** RPE-1 cells were treated with 500 nM reversine for 48 h then with 1 µM AZD1775 for 6 h. **(H)** Immunofluorescence representative images of chromosome spreads showing CENP-A in green and DNA in blue. **(I)** Quantification of the proportion of chromosome spreads from (H) (mean ± S.E.M of N=2; a minimum of 54 spreads were analyzed per group; two-way ANOVA; AZD1775 Cen^OFF^ vs AZD1775 + Reversine Cen^OFF^ ****p<0.0001). Scale bar = 5 µm.

The observation of such a phenotype, characterized by DNA exclusion from the spindle during mitosis, raises the question of the localization of centromere proteins (CENPs) in AZD1775-treated cells. Using anti-centromere antibodies (ACA) staining, we found that WEE1 inhibition caused the detachment of CENPs from the DNA, as evidenced by a chromatin mass located on the outside of the spindle and CENPs clustered on spindle fibers (Figure 3C).

Similar results were obtained using CENP-A and CENP-B as centromeric markers (Figure 3-figure supplement 1A,B). We further distinguished two phenotypes, that we referred to as (i) **‘central chromatin mass’** where the DNA is still mildly attached to the spindle (Figure 3D, central panel) and (ii) **‘side chromatin mass’** where DNA is completely devoid of CENPs that are clustered on the spindle (Figure 3D, right panel). AZD1775-treated control and ATIP3-depleted cells exhibited equal proportions of the ‘central chromatin mass’ phenotype, whereas the ‘side chromatin mass phenotype’ was the major phenotype in ATIP3-depleted cells (Figure 3E). The detachment of DNA from the spindle prompted us to evaluate chromosome integrity by performing chromosome spreads. Following WEE1 inhibition, we observed two distinct types of chromosome spreads. The first type, termed as “Cen^ON^” showed CENPs attached to the chromosomes, although a few acentric chromosomes were present. In the second type, denoted as “Cen^OFF^”, CENPs were clustered and chromosomes were disintegrated, a state that we referred to as chromosome pulverization (Figure 3F, and Figure 3-figure supplement 1C). The latter spreads were largely reminiscent of the ‘side chromatin mass’ phenotype. Importantly, WEE1 inhibition increased the percentage of “Cen^OFF^” spreads in ATIP3-depleted cells (65% compared to 23% in control cells) (Figure 3G).

We examined whether WEE1 inhibition may trigger such chromosome pulverization in aneuploid cells, independently of ATIP3 deficiency. WEE1 inhibition induced chromosome pulverization in RPE-1 cells rendered aneuploid using reversine, but not in their near-diploid counterparts (Figure 3H-I). Similar results were obtained in the chromosomally stable HCT116 cancer cell line, where WEE1 inhibition alone exhibited negligible effects but triggered chromosome pulverization upon increasing aneuploidy using reversine (Figure 3-figure supplement 2A-C). Accordingly, elevating aneuploidy using reversine in HeLa cells, that are already aneuploid (Figure 3-figure supplement 2D), further increased the occurrence of chromosome pulverization (Figure 3-figure supplement 2E,F). These results underscore the crucial relationship between aneuploidy and cell sensitivity to WEE1 inhibition. To further examine centromeric aberrations, we investigated the binding of CENP-B to its specific DNA box, CENP-B box. Immunofluorescence coupled to Fluorescence *in situ* hybridization (FISH) was performed to reveal both CENP-B and CENP-B box. Following WEE1 inhibition, the CENP-B box was no longer detected in “Cen^OFF^” spreads in which the protein CENP-B was clustered (Figure 3-figure supplement 2G,H), indicating loss or disruption of centromeric DNA. This suggests that inhibition of WEE1 results in DNA fragmentation in the centromeric region.

We then investigated the mechanisms leading to aberrant phenotypes in mitosis and chromosome pulverization after exposure to WEE1 inhibition. We hypothesized that chromosome pulverization during mitosis may stem from events that occur earlier in S-phase. WEE1 inhibition was previously shown to cause replication stress by exhausting replication origins, leading to replication fork stalling^24^. Stalled forks can then be recognized by a nuclease for processing^32^. Importantly, uncontrolled action of nucleases can lead to excessive fork degradation and massive DNA double-strand breaks (DSBs). To investigate the early effects of WEE1 inhibition in S-phase before mitotic entry, cells were treated for 2 h with AZD1775 (Figure 4A). WEE1 inhibition did not change the proportion of cells in S-phase, as indicated by the proportion of EdU-positive cells (Figure 4-figure supplement 1A-B). In line with higher DNA replication levels, WEE1 inhibition induced higher levels of EdU incorporation although no differences were observed between control and ATIP3-depleted cells (Figure 4-figure supplement 1C). One of the early events of WEE1 inhibition was the induction of replication stress, as shown by elevated levels of phosphorylated replication protein A (RPA32 pS4/S8) (Figure 4A,B). Interestingly, ATIP3-depleted cells showed significantly higher levels of replication stress in response to WEE1 inhibition than control cells (Figure 4A,B). As excessive replication stress can induce the formation of DSBs^33^, we assessed DNA damage by analyzing pan-nuclear H2AX phosphorylation (γH2AX). DNA damage levels were elevated upon WEE1 inhibition and even further in ATIP3-depleted cells (Figure 4C,D). In line with these results, our *in vivo* studies revealed that ATIP3-deficient tumors exhibit higher levels of DNA damage following short treatment with AZD1775 (Figure 4-figure supplement 1D,E). Inversely, the percentage of EdU-positive cells with 53BP1 foci was reduced upon WEE1 inhibition, suggesting either an impaired DNA damaged response or reduced DNA repair activity (Figure 4-figure supplement 1F,G).

**Figure 4:**
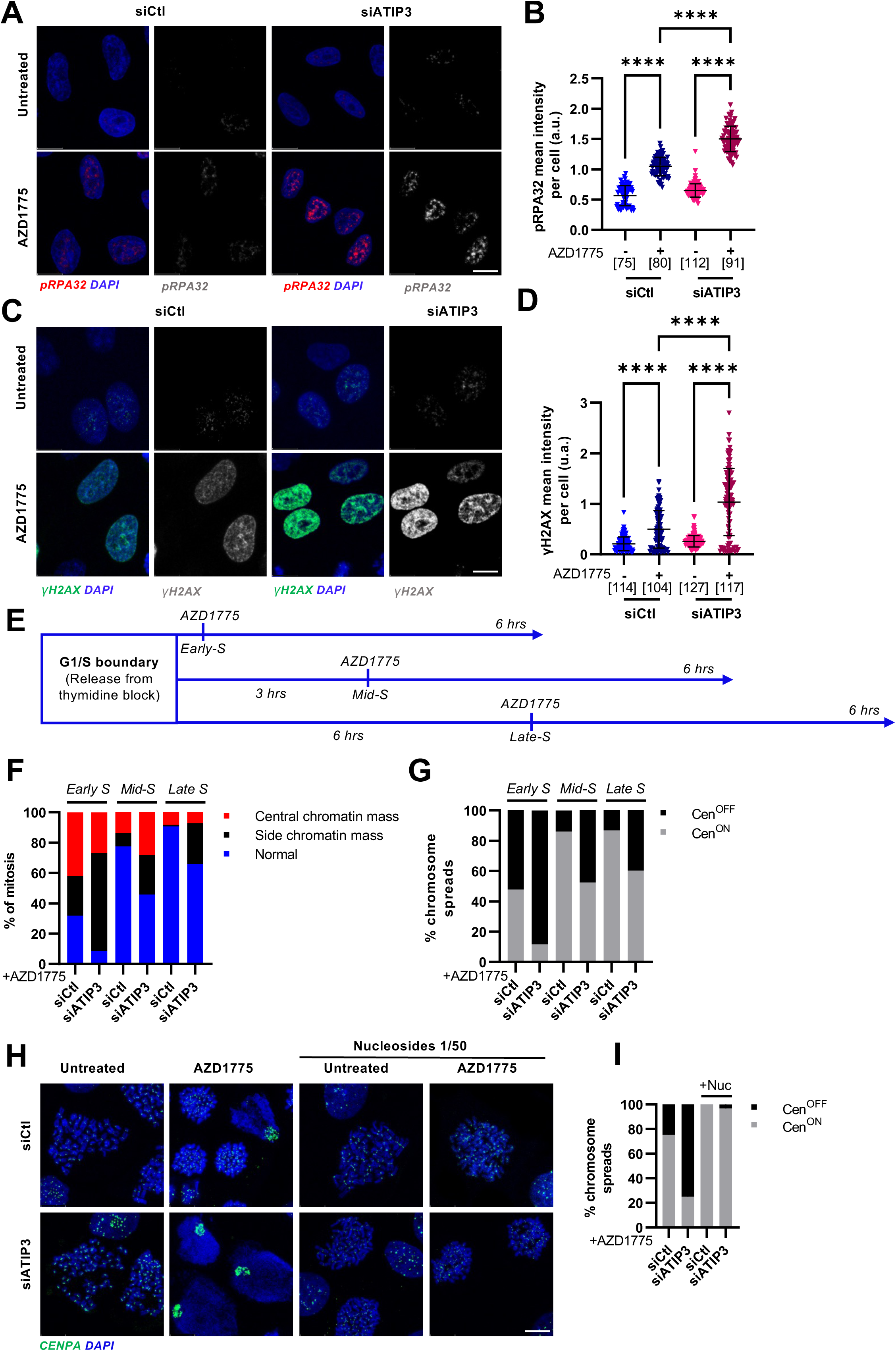
Chromosome pulverization in mitosis is due to heightened levels of replication stress and DNA damage in S-phase. **(A-D)** HeLa cells expressing or not ATIP3 were treated or not with 500 nM AZD1775 for 2 h. **(A)** Immunofluorescence representative images showing phosphorylated RPA32 (S4/S8) in red and DNA in blue. **(B)** Scattered dot plot of phosphorylated RPA32 mean intensity per nucleus (normalized to DAPI). **(C)** Immunofluorescence representative photographs showing γH2AX in green and DNA in blue. **(D)** Scattered dot plot of γH2AX mean intensity per nucleus (normalized to DAPI). **(B-D)** The number of analyzed cells is in brackets (mean ± S.D.; Kruskal-Wallis test followed by Dunn’s multiple comparisons; ****p<0.0001). Scale bar = 20 µm. **(E)** Treatment scheme of HeLa cells synchronized at the G1/S boundary. Cells were either treated with 500 nM AZD1775 for 6 h directly after release from thymidine block (top axis, early-S); after 3 h into S phase progression (middle axis, mid-S), or after 6 h into S phase progression (bottom axis, late-S). **(F)** Quantification of the proportions of the mitotic phenotypes as described in (E) (mean ± SEM. of N=3; a minimum of 133 cells were analyzed per group; two-way ANOVA; side chromatin mass in siCtl early-S *vs*. late-S p<0.0001; side chromatin mass in siATIP3 early-S *vs*. late-S p<0.0001) **(G)** Quantification of the proportions of chromosome spreads as described in (E) (mean ± SEM. of N=3; a minimum of 80 spreads were analyzed per group; two-way ANOVA; Cen^OFF^ in siCtl early-S *vs*. late-S p<0.0001; Cen^OFF^ in siATIP3 early-S *vs*. late-S p<0.0001). **(H)** Immunofluorescence representative images of chromosome spreads from HeLa cells expressing or not ATIP3 and treated with 500 nM AZD1775 for 6 h in the presence or absence of nucleosides (1/50). CENP-A is shown in green and DNA in blue. **(I)** Quantification of the proportion of chromosome spreads shown in (H) (mean ± SEM. of N=2; a minimum of 61 spreads were analyzed per group; two-way ANOVA; Cen^OFF^ in siCtl AZD1775 *vs*. AZD1775 + Nucleosides p<0.0001; Cen^OFF^ in siATIP3 AZD1775 *vs*. siATIP3 AZD1775 + Nucleosides Cen^OFF^ p<0.0001). Scale bar = 5 µm.

Importantly, after 6 h of treatment, when cells progressed to mitosis, the impact of WEE1 inhibition on DNA replication and damage remained pronounced in ATIP3-depleted cells. This is shown by the higher proportion of hyperphosphorylated RPA32 and γH2AX-positive mitoses (Figure 4-figure supplement 2A-D). Of note, EdU-positive mitotic cells also occurred following WEE1 inhibition, and to a greater extent in ATIP3-depleted cells (Figure 4-figure supplement 2C,E), indicating either replication stress-associated mitotic DNA synthesis (MIDAS)^34^ or abrupt mitotic entry of replicating cells. The observed higher levels of damage in ATIP3-depleted mitotic cells led us to investigate whether they could be attributed to the presence of pre-existing replication stress prior to WEE1 inhibition. Cells were pretreated with a low dose of the DNA polymerase inhibitor aphidicolin for 2 h and then exposed for 6 h to AZD1775. When control cells were exposed to aphidicolin and AZD1775, they exhibited the same proportions of hyperphosphorylated RPA32 and γH2AX mitoses as those observed in ATIP3-depleted cells treated with AZD1775 alone (Figure 4-figure supplement 3A-D). These results point towards the existence of low levels of endogenous replication stress or a compromised replication stress response in ATIP3-depleted cells. Indeed, ATIP3-depleted cells showed higher levels of endogenous 53BP1 nuclear bodies (Figure 4-figure supplement 3E-G) a marker of unresolved replication stress transmitted throughout the cell cycle^35^.

We then investigated whether the effects of WEE1 inhibition in S-phase may be responsible for the detachment of CENPs from DNA and the chromosome pulverization observed during mitosis. To this end, G1/S synchronized cells were treated at different time points with AZD1775 following the scheme shown in Figure 4E. Control cells that completed at least half of their S-phase (mid-S and late-S) in the absence of AZD1775 presented much less abnormal phenotypes (side or central chromatin mass phenotypes) in mitosis (Figure 4F) and less chromosome pulverization (Figure 4G) compared to cells that were released into S-phase in the presence of AZD1775. Remarkably, a similar trend was observed in ATIP3-depleted cells. When allowed to progress into S phase before exposure to AZD1775, ATIP3-depleted cells displayed diminished occurrence of abnormal mitosis and chromosome pulverization in subsequent mitosis (Figure 4F,G). These results imply that the effects of the WEE1 inhibitor on DNA replication in S phase are crucial for the development of abnormalities during subsequent mitosis. Notably, allowing cells to partially replicate their DNA helps to alleviate mitotic abnormalities.

We investigated whether the restoration of DNA replication by adding nucleosides, which would counteract the nucleotide depletion caused by WEE1 inhibition^24^, could prevent abnormal mitosis and chromosome pulverization. Simultaneous nucleoside supplementation in AZD1775-treated cells led to a comprehensive rescue of replication stress in S-phase, as well as subsequent abnormal mitoses and chromosome pulverization (Figure 4-figure supplement 3H,I) in both control and ATIP3-depleted cells, highlighting the dysregulation of DNA replication as a driving force behind mitotic aberrations induced by WEE1 inhibition.

Knowing that WEE1 inhibition during S phase results in chromosome pulverization in subsequent mitosis, we explored the potential involvement of key molecular players. Notably, the MUS81 endonuclease emerged as a candidate of interest due to its role in inducing DNA breakage at stalled replication forks after WEE1 inhibition^25^. We co-depleted MUS81 and ATIP3 and studied the extent of replication stress and DNA damage after WEE1 inhibition (Figure 5 and Figure 5-figure supplement 1A). MUS81 silencing in ATIP3-depleted cells led to a significant reduction in the levels of DNA damage (Figure 5A-B) and replication stress (Figure 5-figure supplement 1B,C) induced by WEE1 inhibition. However, MUS81 depletion had no impact on the occurrence of the mitotic phenotypes (Figure 5-figure supplement 1D,E) or chromosome pulverization (Figure 5C-D), regardless of whether in control or ATIP3-depleted cells. This suggests that while MUS81 plays a role during S phase, it does not significantly contribute to abnormal mitosis nor chromosome pulverization, implying the involvement of additional factors. This led us to test the implication of three other nucleases (EXO1, MRE11 and DNA2) which are involved in DNA resection during DNA repair (EXO1), DNA double-strand break sensing and processing (MRE11) and replication fork restart (DNA2)^36–38^. Given our results showing the intricate interplay between replication stress, DNA damage and mitotic processes, we reasoned that these nucleases might play roles in the mitotic phenotype and chromosome pulverization induced by WEE1 inhibition. Depletion of either EXO1 or MRE11 did not significantly influence abnormal mitoses (Figure 5-figure supplement 2A-C) or chromosome pulverization (Figure 5-figure supplement 2D,E), leading us to rule out their direct involvement as primary mediators of these effects. In contrast, DNA2 depletion rescued the abnormal mitotic phenotypes in ATIP3-depleted cells (Figure 5-figure supplement 3) and prevented chromosome pulverization (15% ‘Cen^OFF^’ spreads when DNA2 is co-depleted with ATIP3 *vs*. 55% ‘Cen^OFF^’ spreads in ATIP3 depleted cells) (Figure 5E-F). Furthermore, the depletion of DNA2 prevented the excessive levels of DNA damage induced in ATIP3-deficient cells in response to WEE1 inhibition (Figure 5G-H). These findings point to a pivotal role of DNA2 in orchestrating the observed mitotic phenotypes subsequent to S-phase defects induced by WEE1 inhibition in aneuploid cancer cells.

**Figure 5:**
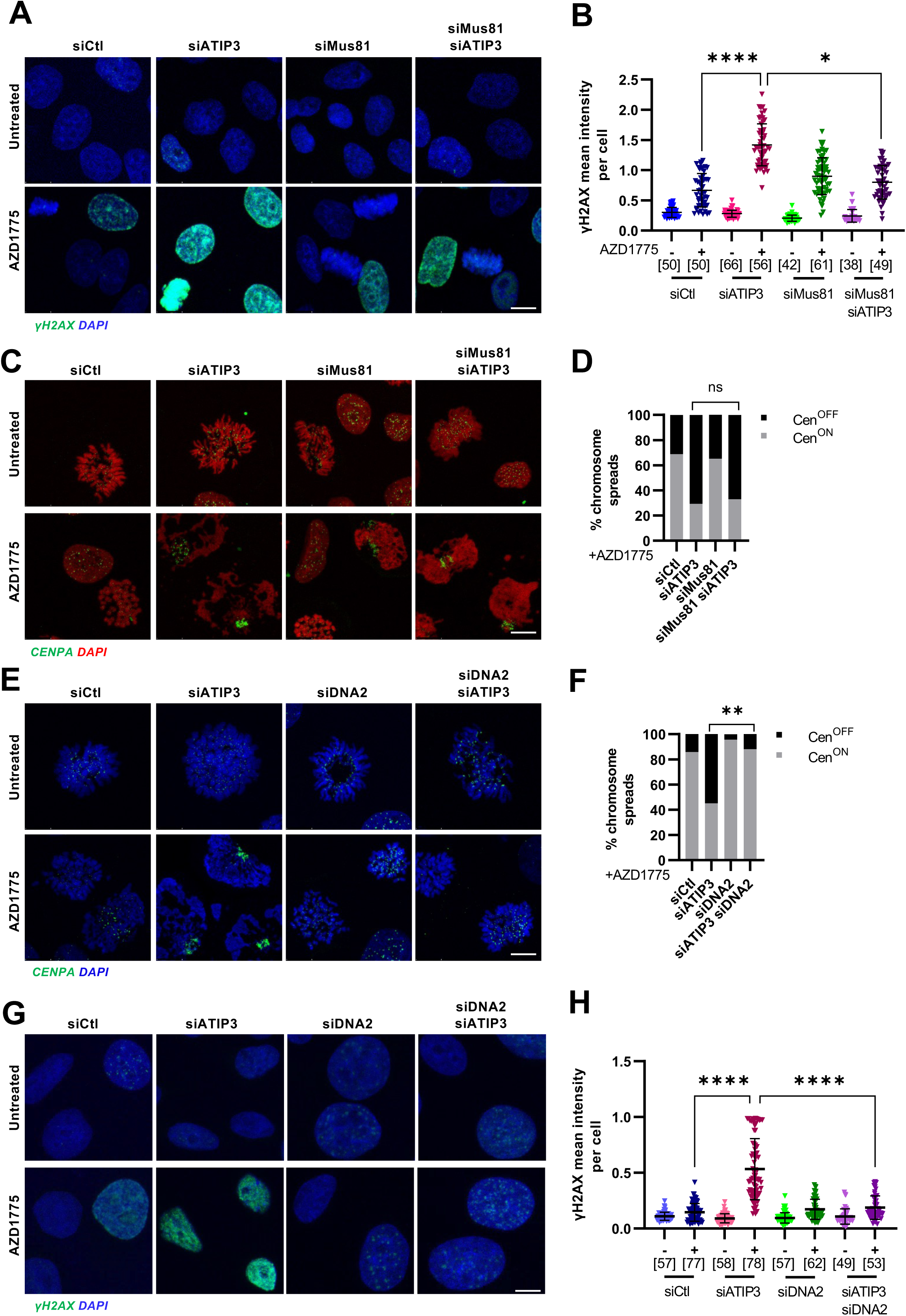
DNA2 nuclease is responsible for chromosome pulverization in mitosis. **(A-D)** HeLa cells transfected with control, ATIP3, MUS81 or a combination of ATIP3 and MUS81 siRNAs were treated or not with 500 nM AZD1775 for 2 h (A-B) or 6 h (C-D). **(A)** Immunofluorescence representative images showing γH2AX in green and DNA in blue. **(B)** Scattered dot plot of γH2AX mean intensity per nucleus (normalized to DAPI). The number of analyzed cells is in brackets (mean ± S.D.; Kruskal-Wallis test followed by Dunn’s multiple comparisons; ****p<0.0001; *<0.05). Scale bar = 20 µm **(C)** Immunofluorescence representative images of chromosome spreads showing CENP-A in green and DNA in red. **(D)** Quantification of the proportion of chromosome spreads shown in (C) (mean ± S.E.M of N=2; a minimum of 67 spreads were analyzed per group; two-way ANOVA showing no significance (p>0.99) between Cen^OFF^ in siATIP3 *vs*. siATIP3 + siMUS81). Scale bar = 5 µm. **(E-H)** HeLa cells transfected with control, ATIP3, DNA2 or a combination or ATIP3 and DNA2 siRNAs and treated or not with 500 nM AZD1775 for 6 h (E-F) or 2 h (G-H). **(E)** Immunofluorescence representative images of chromosome spreads showing CENP-A in green and DNA in blue. **(F)** Quantification of the proportion of chromosomes spreads shown in (E) (mean ± S.E.M of N=2; a minimum of 78 spreads were analyzed per group; two-way ANOVA; Cen^OFF^ in siDNA2 *vs*. siDNA2 + siATIP3 **p<0.01). Scale bar = 5 µm. **(G)** Immunofluorescence representative images of chromosomes spreads showing γH2AX in green and DNA in blue. **(H)** Scattered dot plot of γH2AX mean intensity per nucleus (normalized to DAPI). The number of analyzed cells is in brackets (mean ± S.D.; Kruskal-Wallis test followed by Dunn’s multiple comparisons; ****p<0.0001; *<0.05). Scale bar = 20 µm.

## Discussion

The findings presented in this study provide compelling evidence for increased sensitivity of aneuploid cells to WEE1 inhibition through severe chromosome pulverization. Using ATIP3 depletion as a model to increase aneuploidy levels, we demonstrate heightened susceptibility of highly aneuploid cells to WEE1 inhibition, driven by a combination of processes including increased levels of replication stress and DNA damage in S-phase combined with an accelerated cell cycle progression, ultimately leading to DNA2 nuclease-mediated chromosome pulverization in mitosis and subsequent cell death.

In aneuploid cells, the uneven segregation of chromosomes and compromised DNA replication machinery routinely challenge cellular integrity^27,39^(Figure 6 *left panel*). In these cells, WEE1 inhibition intensifies replication stress and DNA damage to catastrophic levels, culminating in replication failure. In this context, untimely activation of CDK1 by WEE1 inhibition triggers a premature entry into mitosis, which, when combined with the defects in S-phase, causes aberrant mitotic phenotypes, chromosome pulverization and massive cell death (Figure 6 *right panel*).

**Figure 6:**
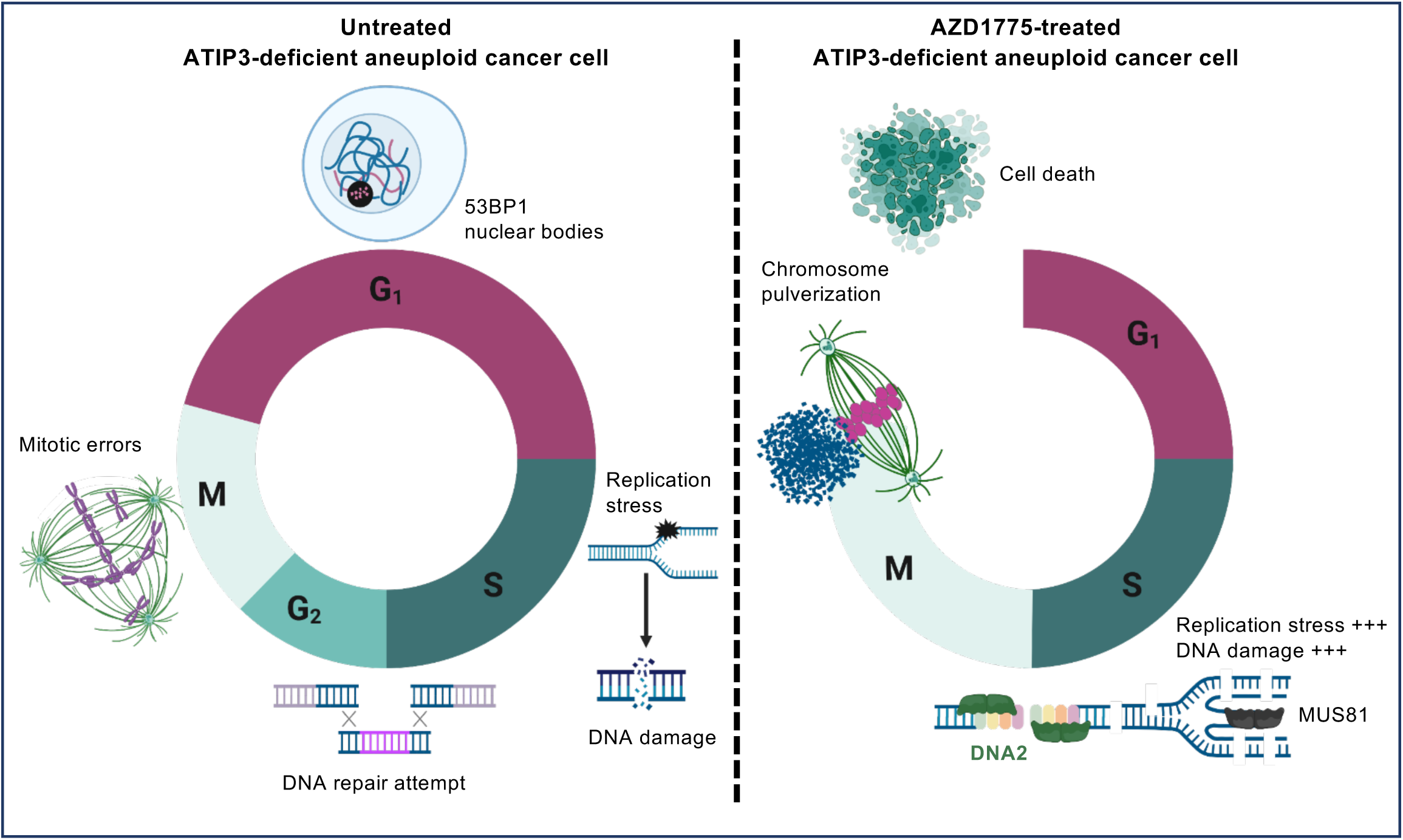
Graphical summary of the proposed working model. ***Left panel:*** During S-phase, aneuploid cells undergo replication stress due to the uneven allocation of chromosomes and flawed DNA replication machinery. This leads to the formation of DNA lesions and stalled replication forks. Consequently, DNA damage response pathways are activated with suboptimal DNA repair mechanisms. As aneuploid cells transition through mitosis, they experience chromosome missegregation causing dissemination of aneuploidy to subsequent daughter cells. ***Right panel:*** WEE1 inhibition in aneuploid cells increases levels of replication stress and DNA DSBs in S-phase through the unrestricted activity of the MUS81 endonuclease and the DNA2 helicase/nuclease. The untimely activation of CDK1 after WEE1 inhibition forces cells to enter mitosis prematurely with under-replicated and damaged DNA. DNA2 helicase/nuclease causes massive chromosome pulverization which leads to massive cell death.

Importantly, the aberrant mitotic phenotypes, characterized by the detachment of centromere proteins from DNA and chromosome pulverization, are due to the effects of WEE1 inhibition in S-phase. Indeed, when cells are exposed to WEE1 inhibition after the completion of S-phase, the occurrence of aberrant mitosis and chromosome pulverization is drastically reduced. Our results favor the hypothesis that replication stress induced by WEE1 inhibition in aneuploid cells reaches intolerable levels that result in excessive nuclease activity during S-phase, leading to massive induction of double-strand breaks. In line with other studies^24,25,40^, we found that the MUS81 endonuclease contributes to DNA damage induced by WEE1 inhibition in aneuploid cells. However, MUS81 depletion was not sufficient to rescue either the mitotic phenotypes or chromosome pulverization, pointing to the existence of additional molecular mechanisms.

In situations where cells undergo premature entry into mitosis while having damaged and under-replicated genomes, one would expect late-replicating regions such as centromeric DNA to remain incompletely replicated despite cell cycle progression. Notably, proteins implicated in DNA repair and replication stress pathways may become enriched during centromeric replication^41^, as these genomic loci are fragile and challenging to replicate. The DNA2 helicase/nuclease, a putative substrate of CDK1^42^, has high affinity for centromeric DNA^43^. Therefore, this enzyme emerges as a pivotal player in the context of centromeric DNA replication to resolve challenging DNA structures. Upon WEE1 inhibition, CDK1 ectopic activation may be responsible for unregulated activity of DNA2. Accordingly, depletion of DNA2 rescues both DNA damage and abnormal mitotic phenotypes, and prevents the severe chromosome pulverization associated with WEE1 inhibition.

Other studies have described a similar aberrant phenotype in response to WEE1 inhibition where cells underwent premature mitosis with under-replicated DNA. This phenotype was referred to as ‘centromere fragmentation’ based on the observation of centromeres being spatially separated from the main mass of chromosomes^44–46^. Our data provide further molecular insights into this phenotype and support a model in which centromere proteins are detached from DNA in a process orchestrated by the action of nucleases. In mitosis, the distinctive “side chromatin mass” phenotype driven by WEE1 inhibition is a result of unregulated DNA2 activity, which breaks centromeric DNA causing the separation of CENPs from the DNA.

This study unveils intricate molecular mechanisms that underlie the consequences of WEE1 inhibition in aneuploid cells, and highlights the DNA2 helicase/nuclease as a new molecular player of cell cycle regulation upon inhibition of WEE1 kinase. Our findings hold significant implications for breast cancer therapies, particularly in revealing the vulnerability of highly aneuploid breast cancers to WEE1-targeted therapies. This is of particular importance in the context of recent clinical trials using WEE1 inhibitors as anti-cancer drugs^47^.

## Materials and methods

### Cell culture, synchronization, and treatment

All cell lines were grown in a sterile cell culture environment and were routinely tested for mycoplasma contamination. MDA-MB-231 (breast cancer cell line), MDA-MB-468 (breast cancer cell line), HeLa (cervical carcinoma cell line) and RPE-1 (hTERT immortalized retinal pigmented cell line) cells were cultured in Dulbecco’s modified Eagle’s medium (DMEM

Gibco™, 61965026). SUM52PE (breast cancer cell line) and HCT116 (colon carcinoma cell line) cells were cultured in RPMI-1640 medium. All cell culture media were supplemented with 10% fetal bovine serum (FBS). MDA-MD-231 shCtl and shATIP3, SUM52PE shCtl and shATIP3, and HeLa H2B-mCherry cells were previously described^17^. Cell lines were grown under standard cell culture conditions in CO_2_ incubators (37□°C; 5% CO_2_). Cells were routinely tested for absence of mycoplasma contamination using Venor® GeM Advance Kit (MB Minerva biolabs®).

For synchronization at the G1/S boundary, HeLa cells were treated with 2□mM thymidine for 18□h. Thymidine was washed out and cells were released into fresh media for 8□h followed by a second exposition to 2□mM thymidine for 18□h.

All drugs were purchased from Selleckchem and resuspended in DMSO. AZD1775 was used at 500 nM for all experiments except for cell viability assessment where it was used at increasing concentrations. RO-3306 was used at 10 µM, aphidicolin at 100 nM and reversine at 500 nM. Media was supplemented with nucleosides (EmbryoMax® Nucleosides, Sigma) 1/50 for DNA replication rescue experiments.

### siRNA transfection

Specific and control scrambled siRNA oligonucleotides were purchased from Dharmacon (Horizon Discovery).

The following sequences were used:

ATIP3 (5’-UGGCAGAGGUUUAAGGUUA-3’);

DNA2: GCUAAACCGUGAAGCAAGA, CUACGUCACUUUAAAGAUG, ACAGUUGCCUGCAUUCUAA, UGAUAUAGAUACCCCAUUA;

EXO1: GAAGUUUCGUUACAUGUGU, GUAAAUGGACCUACUAACA, ACUCGGAUCUCCUAGCUUU, GUUAGCAGCAUUUGGCAUA;

MRE11 GAUGAGAACUCUUGGUUUA, GAAAGGCUCUAUCGAAUGU, GCUAAUGACUCUGAUGAUA, GAGUAUAGAUUUAGCAGAA;

MUS81: GGGAGCACCUGAAUCCUAA, CAGGAGCCAUCAAGAAUAA, GGGUAUACCUGGUGGAAGA, CAGCCCUGGUGGAUCGAUA;

WEE1: AAUAGAACAUCUCGACUUA, AAUAUGAAGUCCCGGUAUA, GAUCAUAUGCUUAUACAGA, CGACAGACUCCUCAAGUGA

All siRNAs (20 nM) were transfected for 48-72 h using Lipofectamine RNAiMAX (Invitrogen). Silencing efficiency was evaluated by qPCR or Western blot.

### Drug screening

SUM52-PE breast cancer cells (2,000) were seeded in round bottom 96-well ultra-low attachment plates (Thermo Fisher) to form MCSs as previously described^17^. MCSs were treated with increasing doses of each kinase inhibitor for 3 days. Viability was determined by measurement of ATP using ATPlite assay (Perkin Elmer). Dose-response curves were generated and fitted with IC50 values using GraphPad Prism (dose-response with variable slope model).

### Xenografts and drug treatment

For tumor formation, 5 million MDA-MB-468 (shCtl or shATIP3) cells were mixed with Geltrex (Gibco) and PBS (1:1) and injected in 100 μl subcutaneously into the left flank of 6–8-week-old NOD SCID gamma (NSG) mice (Charles River). Tumor growth was measured every 4 days using a caliper and volume was assessed as (length × width^2^)/2. When the tumor volumes reached approximately 60-70 mm^3^, mice were randomly segregated into 4 groups (n = 9 per group). Mice were treated daily with vehicle (0.5% methylcellulose) or 90 mg/kg AZD1775 (provided by AstraZeneca) (in 0.5% methylcellulose) via oral gavage for 26 days. Body weight was measured every 4 days as an indicator of toxicity. Animal experiments were performed in accordance with guidelines and approved by the ethical committee of the animal facility of Gustave Roussy Institute, Villejuif, France.

### Live cell imaging

For quantification of mitotic entry, HeLa cells were transfected with control or ATIP3 siRNAs for 24 h then synchronized using double thymidine block. Cells were released in the presence or absence of 500 nM AZD1775 and filmed using an incucyte at a magnification of 10x every 10 minutes for 6 h. For fluorescent live cell imaging, HeLa-mCherryH2B cells were transfected for 48 h with control or ATIP3 siRNAs, then incubated with siR-Tubulin dye (10 nM) prior to treatment with 500 nM AZD1775. Cells were imaged using a confocal laser scanning microscope TCS SP8 MP (Leica) using a dry 40X objective, every 6 min for 48 h. Image analysis was performed using ImageJ software.

### Immunofluorescence

Transfected cells were seeded on coverslips one day before treatment. For the analysis of mitotic phenotypes, cells were fixed with ice-cold methanol for 5 min then washed with PBS. For γH2AX and 53BP1 analysis, cells were fixed with 4% paraformaldehyde for 15 min then washed with PBS and permeabilized with 0.5% Triton X-100 in PBS for 30 min. For the analysis of phosphorylated RPA32, cells were pre-extracted using ice-cold 0.2% Triton X-100 for 1 min then fixed with 4% paraformaldehyde and permeabilized. Coverslips were subsequently blocked with 3% BSA in PBS for 1 h at room temperature and incubated with primary antibodies overnight at 4°C. Coverslips were washed with 3% BSA in PBS and incubated with Alexa Fluor-coupled secondary antibodies for 45 min at room temperature. Coverslips were counterstained with DAPI (1/1000) and mounted using FluorSave reagent (Millipore).

For detection of cells in S-phase, EdU incorporation was analyzed using the Click-iT EdU immunofluorescence kit (Invitrogen) according to the manufacturer’s instructions. Cells were pulse labeled with 10□μM EdU for 30 min, fixed and subjected to the Click-iT reaction. The cells were then processed for immunofluorescence staining as described.

For immunofluorescence in 3D, MCSs were harvested and fixed in 4% paraformaldehyde, 0.5% Triton X-100 in PBS for 45 minutes, blocked in 3% BSA, 0.5% Triton X-100 in PBS overnight at 4°C and incubated 24 h in primary antibodies and overnight in secondary antibodies. MCSs were deposited onto slides and sealed with mounting media and coverslips.

The following primary antibodies were used: rabbit anti-pericentrin (ab4448; Abcam, 1/1000), rat anti-alpha-tubulin (ab6160; Abcam, 1/1000), rabbit anti-CENP-B (ab25734, Abcam, 1/1000), mouse anti-CENP-A (ADI-KAM-CC006-E, ENZO, 1/1000), human ACA (AB_2939058, Antibodies Inc., 1/2000), mouse γH2AX (05-630, Millipore, 1/1000), rabbit anti-53BP1 (ab172580, abcam 1/1000), RPA32 phospho S4/S8 (A300-245A, Bethyl, 1/1000), rabbit anti-phospho-histone H3 (06-570, Millipore, 1/1000). All secondary antibodies (Alexa Fluor dyes) were purchased from Jackson Immunoresearch.

### Chromosome spreads and FISH

Transfected cells were seeded on coverslips at low density (30% confluency) one day before treatment. Colcemid (KaryoMAX, Gibco) was added to culture media 6 h before spreading at a final concentration of 0.1 µg/ml. Media was replaced with KCl buffer (75 mM) for 10 min at room temperature. Coverslips were centrifuged at 1800 rpm for 3 minutes and fixed with 4% paraformaldehyde, blocked with 3% BSA 0,1% Triton X-100 in PBS for 30 min at room temperature then immunofluorescence was performed as previously described. IF-FISH for CENP-B box detection was modified from Chardon *et al.*^48^. Briefly, immunofluorescence was performed and cells were post-fixed in 2% formaldehyde for 10 mins. Cells were fixed in Carnoy’s fixative for 15 min, rinsed in 80% ethanol and air-dried. Coverslips were incubated with CENP-B box probe-Cy3 (PNAbio) 1:1 in hybridization buffer (20mM Tris, pH 7.4, 60% formamide, 0.5% of blocking reagent (Roche 11096176001)). Samples and probes were denatured by heating on 75°C for 2 mins and incubated at 37°C overnight. Coverslips were washed with 0.4X SSC, counterstained with DAPI (1/1000) and mounted using FluorSave reagent (Millipore).

### Image acquisition and analysis

Images were acquired with a confocal laser scanning microscope Dmi8-SP8 using a 63X objective. 5 to 10 fields were imaged/treatment. The pinhole diameter was set at 1 airy unit for all channels, and the exposure gain for each channel was kept constant in between image acquisition of all samples. Z-stack projection and image analysis was done using LAS-X analysis software. For intensity analysis (phosphorylated RPA32, γH2AX, EdU and DAPI), background was subtracted and areas of interest (nuclei) were delineated by ROIs. Mean intensities of each channel were calculated as the arithmetic mean of the determined gray-scale values in each ROI then normalized to DAPI mean intensities in the same ROI.

### Quantitative PCR

RNA extraction and reverse transcription were performed as previously described^17^. Briefly, RNA was isolated using TRIzol reagent (Thermo Fisher). RNA concentration was assessed using a NanoDrop and Complementary DNA (cDNA) was generated using Superscript II reverse transcriptase (Thermo Fisher) following manufacturer’s protocol.

Quantitative real-time PCR was performed on a Viia7 real-time PCR machine (Thermo Fisher). Gene expression was normalized to RPL13. The following oligonucleotides were used for assessment of gene silencing efficiency:

ATIP3 (F: GGCGGAACAGTGACAATA; R: GCAAATTCACCCATGACGA);

DNA2 (F: GATTTCTGGCACCAGCATAGCC; R: ACACCTCATGGAGAACCGTACC);

EXO1 (F: TCGGATCTCCTAGCTTTTGGCTG; R: AGCTGTCTGCACATTCCTAGCC);

MRE11 (F: CAGCAACCAACAAAGGAAGAGGC; R: GAGTTCCTGCTACGGGTAGAAG);

MUS81 (F: GATCCTACAGCACTTCGGAGAC; R: AAGAGTCCTGGACTTCCGCAAG);

WEE1 (F: GATGTGCGACAGACTCCTCAAG; R: CTGGCTTCCATGTCTTCACCAC).

### Western blotting

Cells were lysed using RIPA buffer containing a cocktail of protease and phosphatase inhibitors. Protein lysates were denatured in Laemmli buffer, separated by SDS–PAGE and transferred onto a PVDF membrane. After blocking with 5% BSA in TBS-0.1% Tween-20, membranes were incubated with the following primary antibodies: rabbit anti-ATIP3 (Aviva ARP44419-P050, 1/1000), anti-phospho-histone H3 (06-570; Millipore, 1/1000), anti-phospho-tyrosine 15 Cdk1 (Cell Signaling Technology; 9111; 1:1,000 dilution), anti-CDK1 phosphorylated substrates (Cell signaling, 1/1000), rat anti-alpha-tubulin (ab6160; Abcam, 1/20000), goat anti-GAPDH (Novus biologicals, 1/50000). Proteins were visualized using horseradish-peroxidase-conjugated secondary antibodies followed by chemiluminescence detection with ECL (Clarity Western ECL substrate, Bio-Rad).

### Statistical analysis

All statistical analysis was performed using GraphPad Prism 9 software. Statistical significance between two groups was determined using two-tailed t-test. When more than two groups were compared, statistical significance was determined using one or two-way ANOVA or Kruskal-Wallis when data distribution did not pass normality tests. P-values < 0.05 were considered significant.

## Acknowledgements

This work has benefited from the facilities and expertise of PETRA (Nicolas Signolle), PFIC (Tudor Manoliu) and PFEP (Mélanie Polrot) platforms of the UMS AMMICa (CNRS UMS 3655 / Inserm US 23) - Gustave Roussy Cancer Campus Villejuif, France. Authors acknowledge help from Dr Daniele Fachinetti (Institut Curie, Paris, France) and Dr Michele Debatisse (Gustave Roussy, Villejuif, France) for providing reagents and Marianne Oulhen (Gustave Roussy, Villejuif, France) for technical assistance. We thank Dr Sophie Polo (Institut d’Epigénétique, Paris, France), Dr Daniele Fachinetti and Dr Renata Basto (Institut Curie, Paris, France), Dr Olivier Gavet and Dr Valeria Naim (Gustave Roussy, Villejuif, France) for constructive discussion throughout the project.

