## Supplementary material for "WEE1 kinase inhibition triggers severe chromosome pulverization in aneuploid cells": Legend to supplemental Figures

**Legends to Figure Supplements**

**Figure 1 - figure supplement 1**. **(A)** Western blot of ATIP3 protein levels in SUM52PE shCtl or shATIP3 MCSs. Loading control: β-tubulin **(B)** Dose-response curves of the 6 differential inhibitors in SUM52PE shCtl or shATIP3 MCSs upon 72 h of treatment. **(C)** IC50 values in µM of each differential inhibitor in SUM52PE shCtl or shATIP3 MCSs upon 72 h of treatment. **(D)** *Top panels*: Dose-response curves of AZD1775 in MDA-MB-468 or MDA-MB-231 shCtl or shATIP3 MCSs upon 72 h of treatment. *Bottom panels*: Western blot of ATIP3 protein levels in shCtl or shATIP3 MCSs. Loading control: GAPDH. **(E)** Plot representing viability of in SUM52PE shCtl or shATIP3 MCSs upon 72 h of treatment with increasing doses of PD0166285. **(F)** Western blot analysis of CDK1 Tyr15 phosphorylation in SUM52PE shCtl or shATIP3 MCS treated or not with 500 nM AZD1775 for 6 h. Loading control: β-tubulin. **(G-H)** RPE-1 cells were treated for 48 h with 500 nM reversine. Reversine was washed out and cells were treated with 10 µM AZD1775 for 72 h before cell viability was assessed. **(G)** Treatment scheme of RPE-1 cells. **(H)** *Left panel*: Violin plot representing chromosome counts. The number of analyzed spreads is indicated between brackets. *Right panel:* Frequency distribution of chromosome counts.

**Figure 2 - figure supplement 1**. **(A-C)** HeLa cells transfected with control or ATIP3 siRNAs and treated with AZD1775 or PD0166285 for 6 h. **(A)** Western blot of ATIP3 protein levels. Loading control: GAPDH. **(B)** Western blot analysis for histone H3 Ser10 phosphorylation. Loading control: β-tubulin. **(C)** Western blot analysis of CDK1 phosphorylated substrates (pTPXK). Increased phosphorylated substrates are indicated in red. Loading control: Actin. **(D)** Immunofluorescence representative images of SUM52PE shCtl or shATIP3 MCSs treated with 500 nM AZD1775 for 14 h showing phosphorylated histone H3 in magenta and DNA in blue. Scale bar = 100 µm. **(E)** Quantification of the percentage of phospho-histone H3 positive cells in MCS shown in (D) (mean ± S.D.; Kruskal-Wallis test followed by Dunn’s multiple comparisons; **p<0.01, ***p<0.001). **(F)** Percentage of cells exhibiting back and forth movement of their DNA around the spindle in mitosis. The number of analyzed cells in indicated between brackets. **(G)** Immunofluorescence representative images of SUM52PE shCtl or shATIP3 MCSs treated with 500 nM AZD1775 for 14 h showing phosphorylated histone H3 in magenta, microtubules in green and DNA in blue. White arrows indicate the exclusion of DNA from the mitotic spindle. Scale bar = 10 µm. **(H)** HeLa cells transfected with control, WEE1, ATIP3 or both siRNAs. *Top panel*: Immunofluorescence representative images showing centrosomes in green, microtubules in red and DNA in blue. *Bottom panel:* quantification of abnormal mitosis (mean ± SEM. of N=3; a minimum of 97 cells were analyzed per group; two-tailed t-test; **p<0.01). **(I-J)** HeLa cells transfected with control or ATIP3 siRNAs were treated with 500 nM AZD1775, 10 µM RO-3306 or a combination of both drugs for 6 h. **(I)** Immunofluorescence representative images showing microtubules in green and DNA in blue. **(J)** Quantification of abnormal mitosis shown in (C) (mean ± SEM. of N=3; the number of analyzed cells is in brackets; one-way ANOVA; ****p<0.0001). Scale bar = 20 µm. **(K)** HeLa cells transfected with control, CDK2, ATIP3 or both siRNAs were treated with 500 nM AZD1775 for 6 h. The plot shows the quantification of abnormal mitosis (mean ± SEM. of N=2; the number of analyzed cells is in brackets; one-way ANOVA; ****p<0.0001)

**Figure 3 - figure supplement 1.** HeLa cells transfected with control or ATIP3 siRNAs were treated with 500 nM AZD1775 for 6 h. **(A)** Immunofluorescence representative images showing CENP-B in magenta, microtubules in green and DNA in blue. **(B)** Immunofluorescence representative images showing CENP-A in green, microtubules in red and DNA in blue. **(C)** Immunofluorescence representative images of chromosome spreads showing CENP-B in magenta and DNA in blue. Scale bar = 20 µm.

**Figure 3 - figure supplement 2. (A-C)** HCT116 cells were treated for 48 h with 500 nM reversine. Reversine was washed out and cells were treated with 500 nM AZD1775 for 6 h before chromosome spreading. **(A)** *Left panel:* Violin plot representing chromosome counts. The number of analyzed spreads is indicated between brackets. *Right panel:* Frequency distribution of chromosome counts. **(B)** Representative images of chromosome spreads showing CENP-A in green and DNA in blue. **(C)** Quantification of the proportion of chromosome spreads shown in (C) (mean ± S.E.M of N=2; a minimum of 53 spreads were analyzed per group) two-way ANOVA; AZD1775 Cen^OFF^ vs AZD1775 + Reversine Cen^OFF^ ****p<0.0001). **(D-F)** HeLa cells were treated for 48 h with 500 nM reversine. Reversine was washed out and cells were treated with 500 nM AZD1775 for 6 h before chromosome spreading. **(D)** *Left panel:* Violin plot representing chromosome counts. The number of analyzed spreads is indicated between brackets. *Right panel:* Frequency distribution of chromosome counts. **(E)** Representative images of chromosome spreads showing CENP-A in green and DNA in blue. **(F)** Quantification of the proportion of chromosome spreads shown in (E) (mean ± S.E.M of N=2; a minimum of 57 spreads were analyzed per group) two-way ANOVA; AZD1775 Cen^OFF^ vs AZD1775 + Reversine Cen^OFF^ ***p<0.001). Scale bar = 5 µm. **(G-H)** HeLa cells transfected with control or ATIP3 siRNAs were treated with 500 nM AZD1775 for 6 h. **(G)** Immunofluorescence representative images of IF-FISH showing CENP-B box in magenta, CENP-B in green and DNA in blue. **(H)** Quantification of the proportion of chromosome spreads shown in (G) (mean ± SEM. of N=2; the number of analyzed spreads is in brackets; two-way ANOVA; Cen^OFF^ in siCtl AZD1775 *vs*. siATIP3 AZD1775 ****p<0.0001). Scale bar = 5 µm.

**Figure 4 - figure supplement 1.** HeLa cells transfected with control or ATIP3 siRNAs were treated with 500 nM AZD1775 for 2 h (A-C). **(A)** Immunofluorescence representative images showing EdU in yellow and DNA in blue. The gray channel representing EdU is shown on the right to each condition. **(B)** Quantification of the percentage of EdU-positive cells shown in (A) (mean ± SEM. of N=3; a minimum of 73 cells were analyzed per group; one-way ANOVA; *ns*). **(C)** Quantification of EdU mean intensity per nucleus (normalized to DAPI) (mean ± S.D.; the number of analyzed cells is in brackets; Kruskal-Wallis test followed by Dunn’s multiple comparisons; ***p<0.001, ****p<0.0001). **(D-E)** Mice xenografted with MDA-MB-468 shCtl or shATIP3 were treated or not with 90 mg/kg of AZD1775 by oral gavage daily for 4 days (1 mice per group) **(D)** Immunohistochemistry representative images showing γH2AX staining in xenografts. **(E)** Quantification of the percentage of γH2AX-positive nuclei shown in (D). **(F-G)** HeLa cells transfected with control or ATIP3 siRNAs were treated with 500 nM AZD1775 for 2 h. **(F)** Immunofluorescence representative images showing 53BP1 in magenta, EdU in yellow and DNA in blue. The gray channel representing 53BP1 is shown on the right to each condition. **(G)** Quantification of the percentage of EdU-positive cells with > 5 53BP1 foci shown in (F) (mean ± SEM. of N=4, a minimum of 173 cells were analyzed per group; one-way ANOVA; ***p<0.001, ****p<0.0001). Scale bar = 20 µm.

**Figure 4 - figure supplement 2.** HeLa cells transfected with control or ATIP3 siRNAs were treated with 500 nM AZD1775 for 6 h. **(A)** Immunofluorescence representative images showing phosphorylated RPA32 (S4/S8) in magenta, EdU in yellow and DNA in blue. The gray channels representing EdU or pRPA32 are shown on the right to each condition. **(B)** Quantification of the percentage of pRPA32-positive mitosis shown in (A) (mean ± SEM. of N=3; a minimum of 42 cells were analyzed per group; one-way ANOVA; *ns*, **p<0.01). **(C)** Immunofluorescence representative images showing γH2AX in red, EdU in green and DNA in blue. The gray channels representing EdU or γH2AX are shown on the right to each condition. **(D-E)** Quantification of the percentage of γH2AX-positive mitosis (D) and EdU-positive mitosis (E) shown in (C) (mean ± SEM. of N=4; a minimum of 50 cells were analyzed per group; one-way ANOVA; *ns*, *p<0.05, **p<0.01, ****p<0.0001). Scale bar = 20 µm.

**Figure 4 - figure supplement 3. (A-D)** HeLa cells transfected with control or ATIP3 siRNAs were pretreated with 100 nM Aphidicolin for 2 h then with 500 nM AZD1775 for 6 h. **(A)** Immunofluorescence representative images showing phosphorylated RPA32 (S4/S8) in magenta and DNA in blue. **(B)** Quantification of the percentage of pRPA32-positive mitosis shown in (A) (mean ± SEM. of N=2; a minimum of 51 cells were analyzed per group; one-way ANOVA; *ns*, ***p<0.001). **(C)** Immunofluorescence representative images showing γH2AX in green and DNA in blue. **(D)** Quantification of the percentage of γH2AX-positive mitosis shown in (C) (mean ± SEM. of N=2; a minimum of 58 cells were analyzed per group; one-way ANOVA; *ns*, ***p<0.001). **(E)** *Left panel:* Immunofluorescence representative images of HeLa cells transfected with control or ATIP3 siRNAs showing 53BP1 nuclear bodies in red, EdU in green and DNA in blue. *Right Panel:* Quantification of the percentage of G1 cells with 53BP1 nuclear bodies (mean ± SEM. of N=5; a minimum of 343 cells were analyzed per group; two-tailed t-test; *ns*, **p<0.01). **(F-G)** HCT116 cells were transfected with control or ATIP3 siRNAs. **(F)** *Right panel:* Western blot of ATIP3 protein levels. Loading control: GAPDH. *Left panel:* Immunofluorescence representative images showing 53BP1 nuclear bodies in magenta, EdU in green and DNA in blue. **(G)** Quantification of the percentage of G1 cells with 53BP1 nuclear bodies (mean ± SEM. of N=4; a minimum of 120 cells were analyzed per group; two-tailed t-test; *ns*, **p<0.01). Scale bar = 20 µm. **(H-I)** HeLa cells transfected with control or ATIP3 siRNAs were simultaneously with nucleosides (1/50) and 500 nM AZD1775 for 2 h (H) or 6 h (I). **(H)** Scattered dot-plot of phosphorylated RPA32 (S4/S8) per nucleus (normalized to DAPI). The number of analyzed cells is in brackets (mean ± S.D.; Kruskal-Wallis test followed by Dunn’s multiple comparisons; ***p<0.001, ****p<0.0001). **(I)** Graph showing the % of mitotic phenotypes (mean ± SEM. of N=1; a minimum of 43 cells were analyzed per group; Side chromatin mass in siATIP3 AZD1775 *vs*. siATIP3 AZD1775 + Nuc p<0.0001).

**Figure 5 - figure supplement 1.** HeLa cells transfected with control, MUS81, ATIP3 or both siRNAs were treated with 500 nM AZD1775. **(A)** qPCR analysis of gene silencing efficiency. **(B)** Immunofluorescence representative images after 2 h of treatment showing phosphorylated RPA32 (S4/S8) in magenta and DNA in blue. **(C)** Quantification of phosphorylated RPA32 mean intensity per nucleus (normalized to DAPI) (mean ± S.D.; the number of analyzed cells is in brackets; Kruskal-Wallis test followed by Dunn’s multiple comparisons; ***p<0.001, ****p<0.0001). Scale bar = 20 µm. **(D)** Immunofluorescence representative images after 6 h of treatment showing CENPs in magenta, microtubules in green and DNA in blue. **(E)** Quantification of the proportions of mitotic phenotypes shown in (D) (mean ± SEM. of N=2; a minimum of 68 cells were analyzed per group; two-way ANOVA; Side chromatin mass in siATIP3 AZD1775 *vs*. siATIP3 + siMUS81 AZD1775 *ns*). Scale bar = 5 µm.

**Figure 5 - figure supplement 2.** HeLa cells transfected with control, EXO1, MRE11, ATIP3 or a combination of siRNAs were treated with 500 nM AZD1775 for 6 h. **(A)** qPCR analysis of gene silencing efficiency. **(B)** Immunofluorescence representative images showing CENPs in green, microtubules in red and DNA in blue. **(C)** Quantification of the proportion of mitotic phenotypes shown in (B) (mean ± SEM. of N=1; a minimum of 67 cells were analyzed per group; two-way ANOVA; Side chromatin mass in siATIP3 AZD1775 *vs*. siATIP3 siMRE11 AZD1775 *ns;* Side chromatin mass in siATIP3 AZD1775 *vs*. siATIP3 siEXO1 AZD1775 *ns*). **(D)** Immunofluorescence representative images of chromosome spreads showing CENP-A in green and DNA in blue. **(E)** Quantification of the proportions of chromosome spreads shown in (D) (mean ± SEM. of N=1; a minimum of 73 spreads were analyzed per group; two-way ANOVA; Cen^OFF^ in siATIP3 AZD1775 *vs*. siATIP3 siMRE11 AZD1775 *ns;* Cen^OFF^ in siATIP3 AZD1775 *vs*. siATIP3 siEXO1 AZD1775 *ns*). Scale bar = 20 µm.

**Figure 5 - figure supplement 3.** HeLa cells transfected with control, ATIP3, DNA2 or a combination or ATIP3 and DNA2 siRNAs and treated with 500 nM AZD1775 for 6 h. **(A)** qPCR analysis of gene silencing efficiency. **(B)** Immunofluorescence representative images of abnormal mitoses showing CENPs in magenta, microtubules in green and DNA in blue. **(C)** Quantification of the proportions of the mitotic phenotypes shown in (A) (mean ± S.E.M of N=2; a minimum of 88 cells were analyzed per group; two-way ANOVA; normal mitoses in siATIP3 *vs*. siATIP3 + siDNA2 **p<0.01). Scale bar = 20 µm.

**Movie S1: Cell division of untreated control HeLa mCherry-H2B cells.** Microtubules are shown in Cyan (siR-tubulin) and the nucleus is in red.

**Movie S2: Cell division of untreated ATIP3-depleted HeLa mCherry-H2B cells.** Microtubules are shown in Cyan (siR-tubulin) and the nucleus is in red.

**Movie S3: Cell division of AZD1775-treated control HeLa mCherry-H2B cells.** Microtubules are shown in Cyan (siR-tubulin) and the nucleus is in red.

**Movie S4: Cell division of AZD1775-treated ATIP3-depleted HeLa mCherry-H2B cells.** Microtubules are shown in Cyan (siR-tubulin) and the nucleus is in red.
