## Supplementary figures and images for "WEE1 kinase inhibition triggers severe chromosome pulverization in aneuploid cells"

### Supplemental Figure 1-1

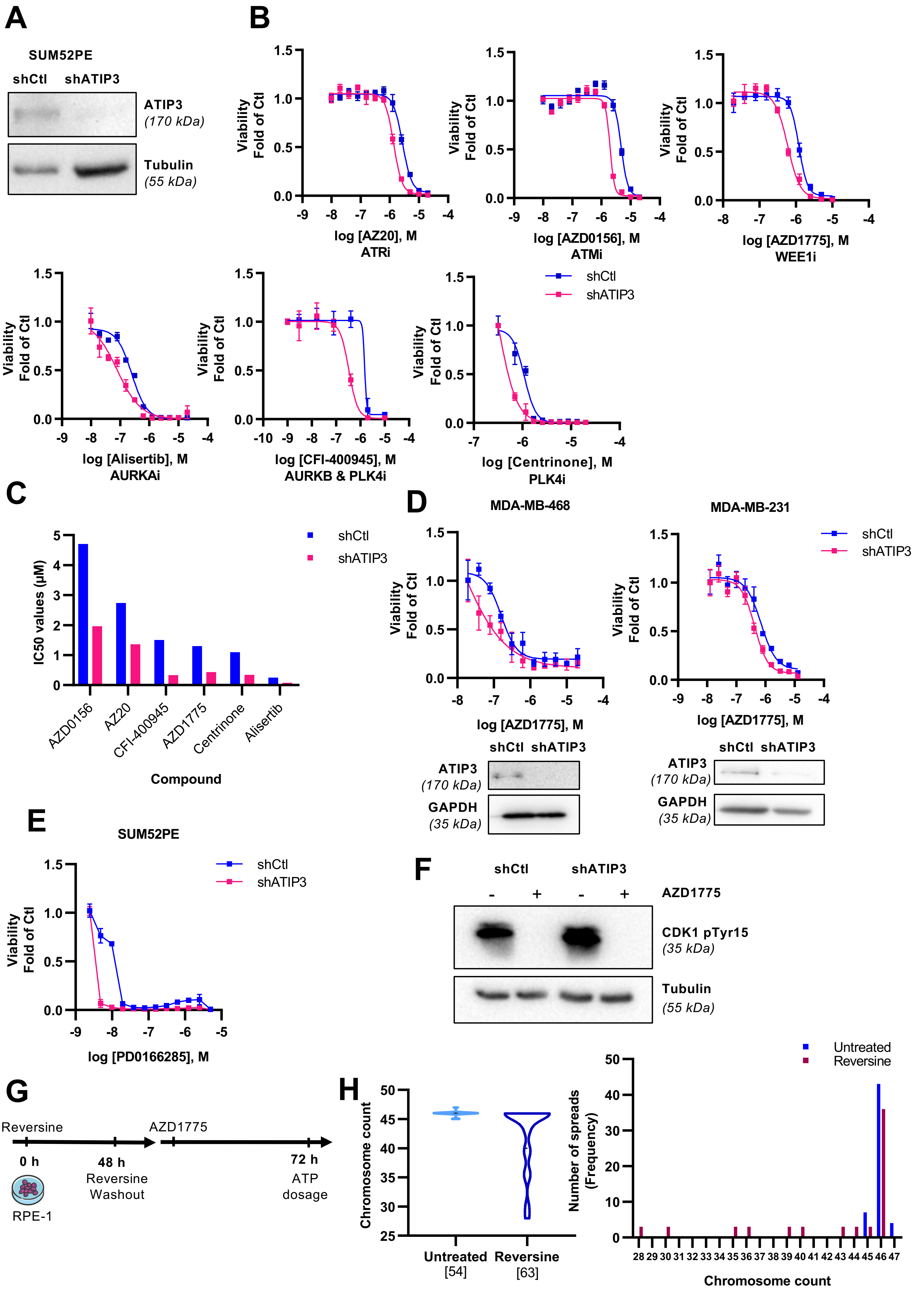


**Figure 1 – figure supplement 1**

### Supplemental Figure 2-1

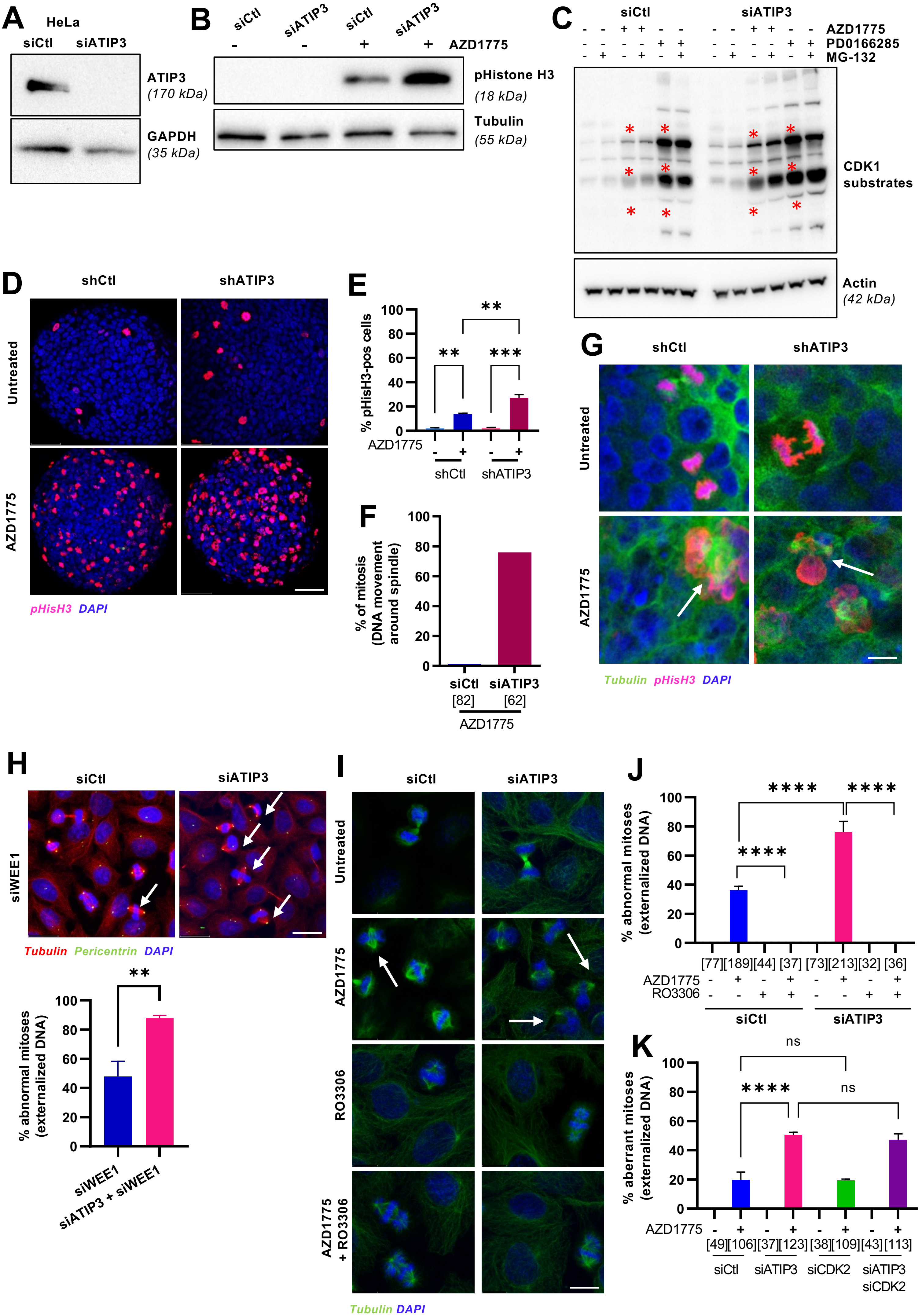


**Figure 2 – figure supplement 1**

### Supplemental Figure 3-1

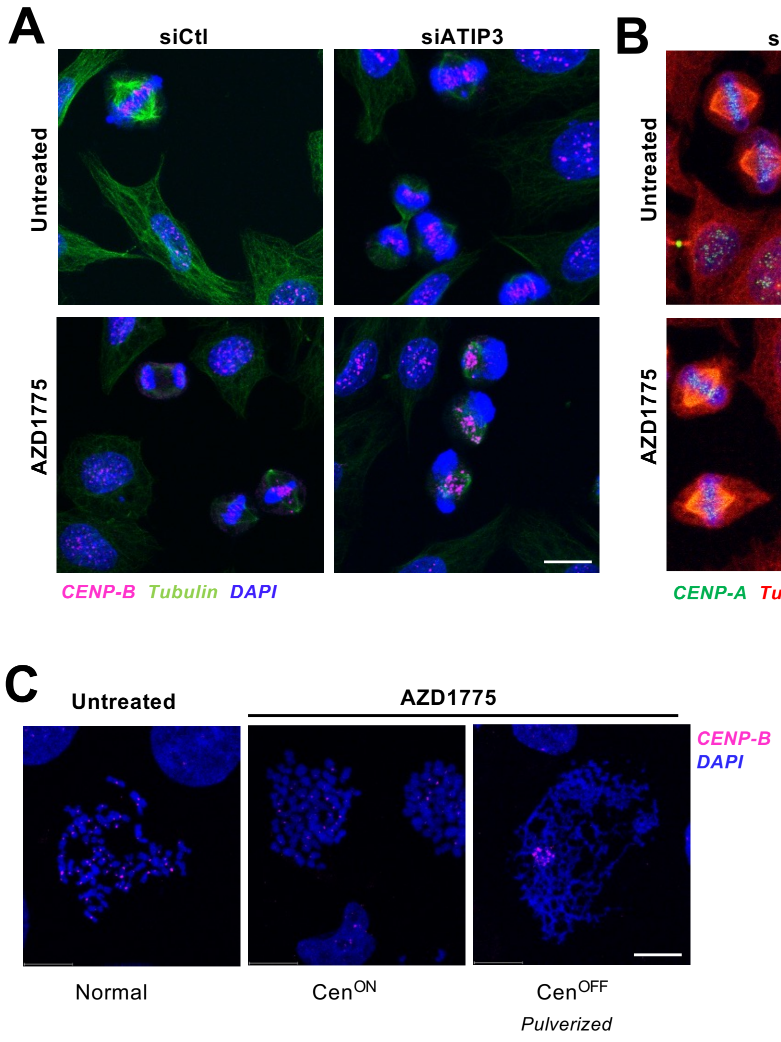


**Figure 3 – figure supplement 1**

### Supplemental Figure 3-2

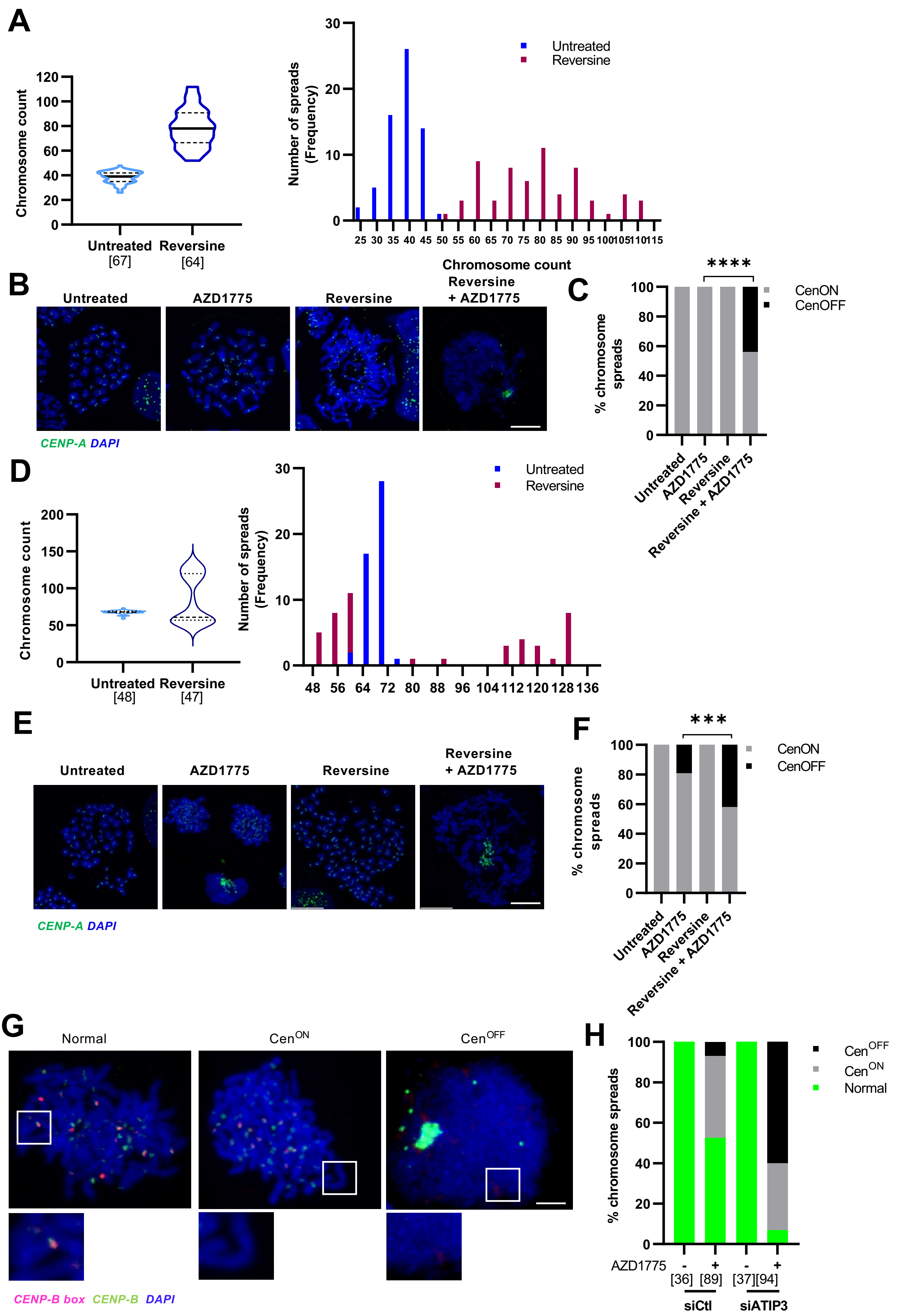


Figure 3 – figure supplement 2

### Supplemental Figure 4-1

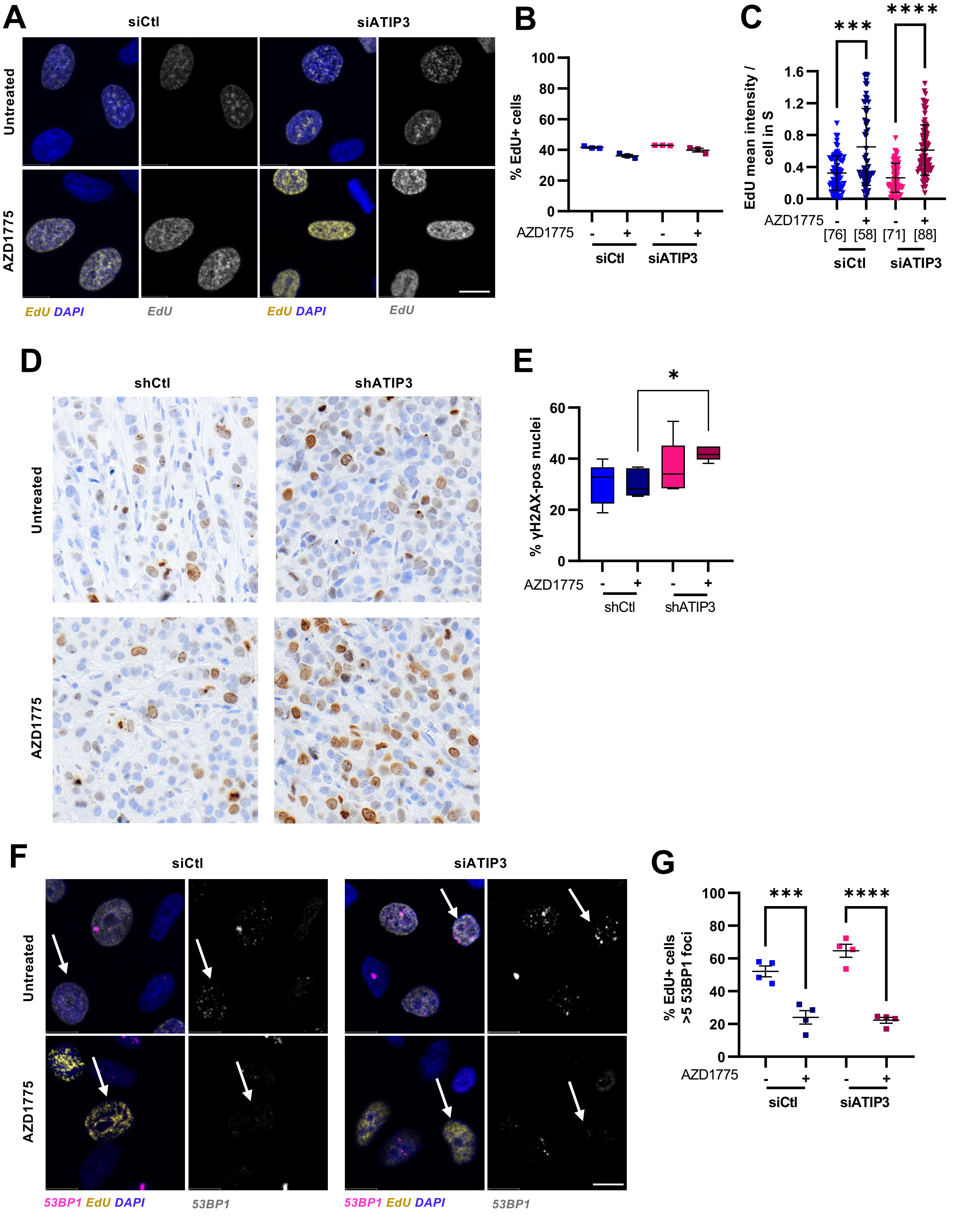


Figure 4 – figure supplement 1

### Supplemental Figure 4-2

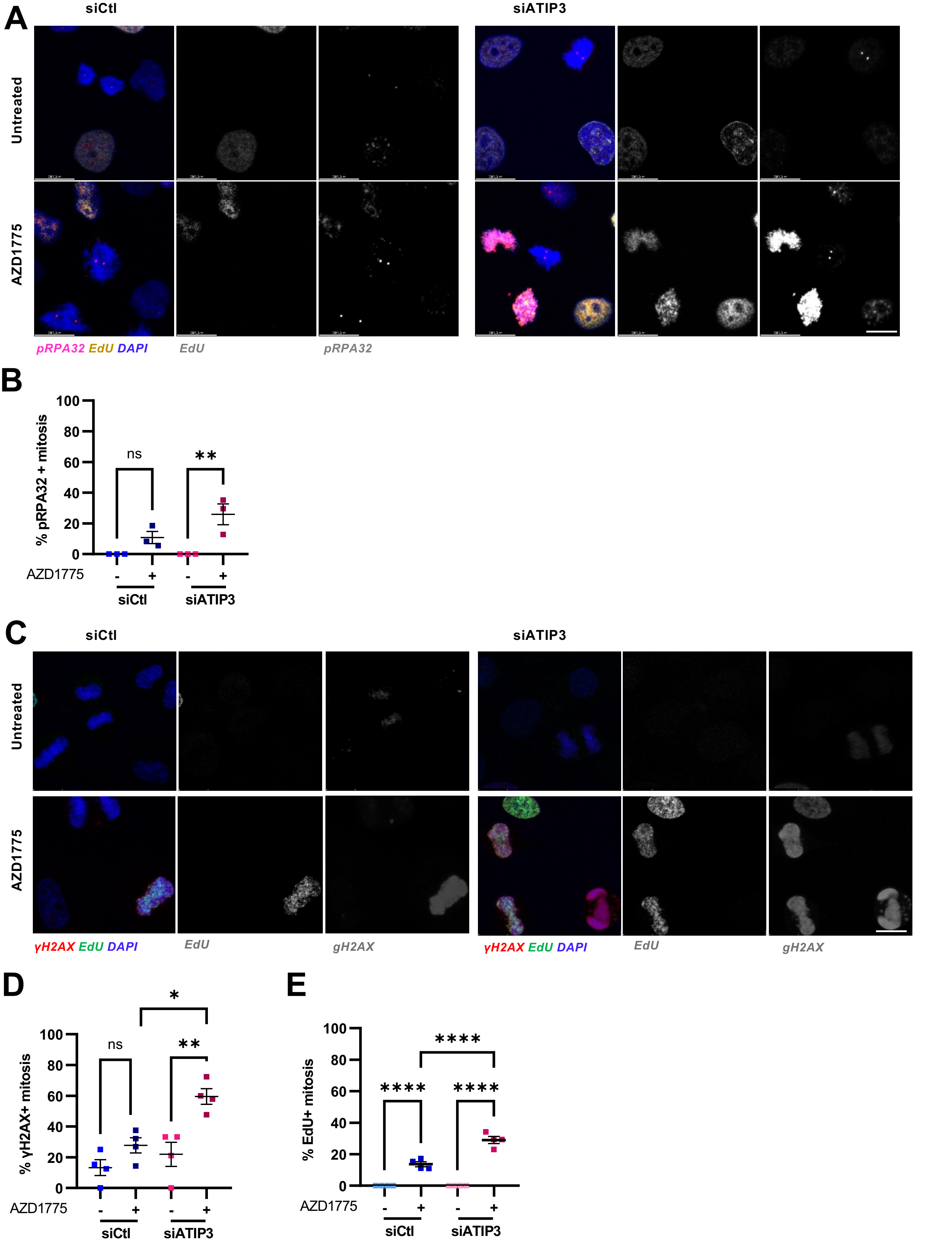


Figure 4 – figure supplement 2

### Supplemental Figure 4-3

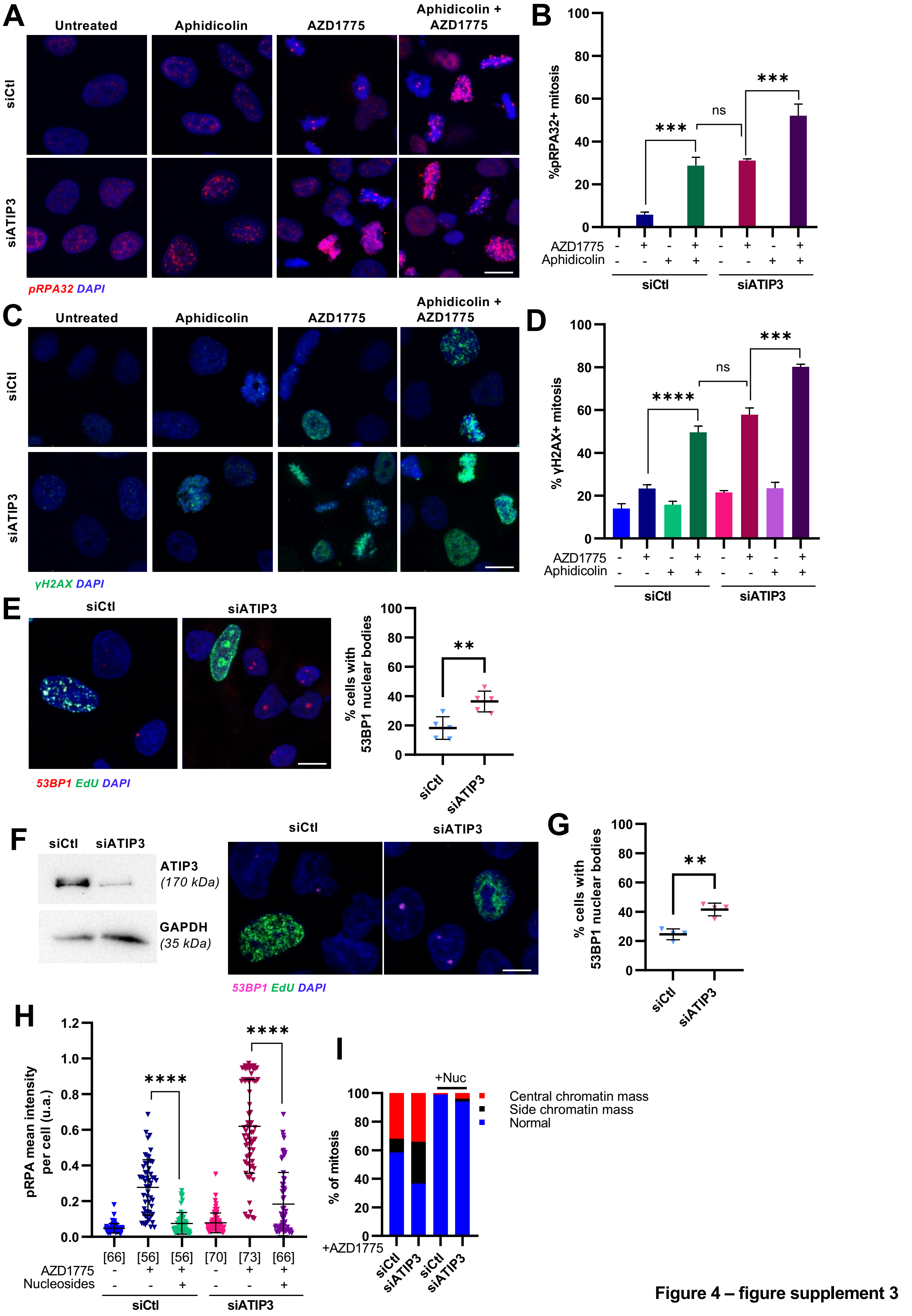


Figure 4 – figure supplement 3

### Supplemental Figure 5-1

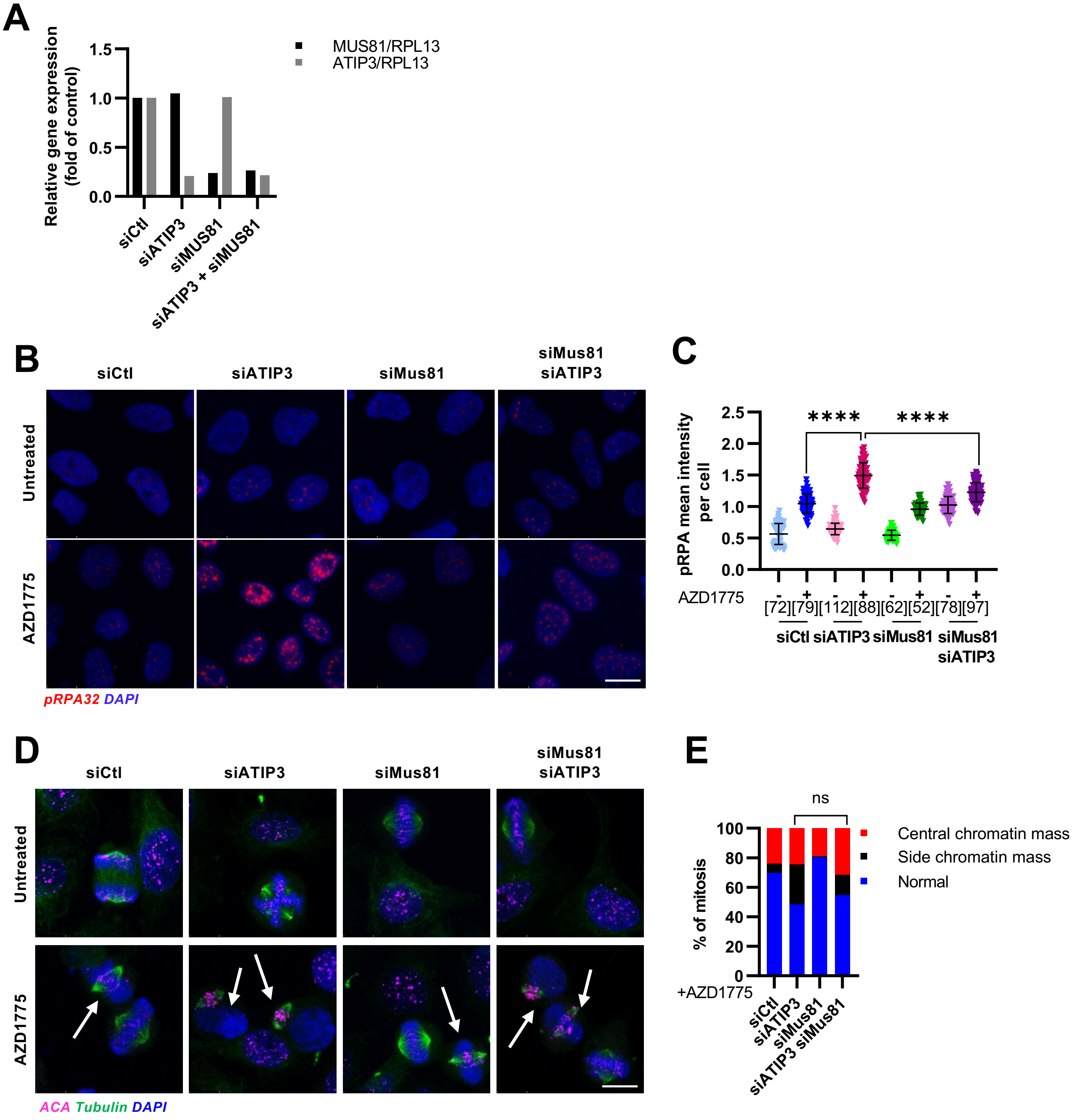


Figure 5 – figure supplement 1

### Supplemental Figure 5-2

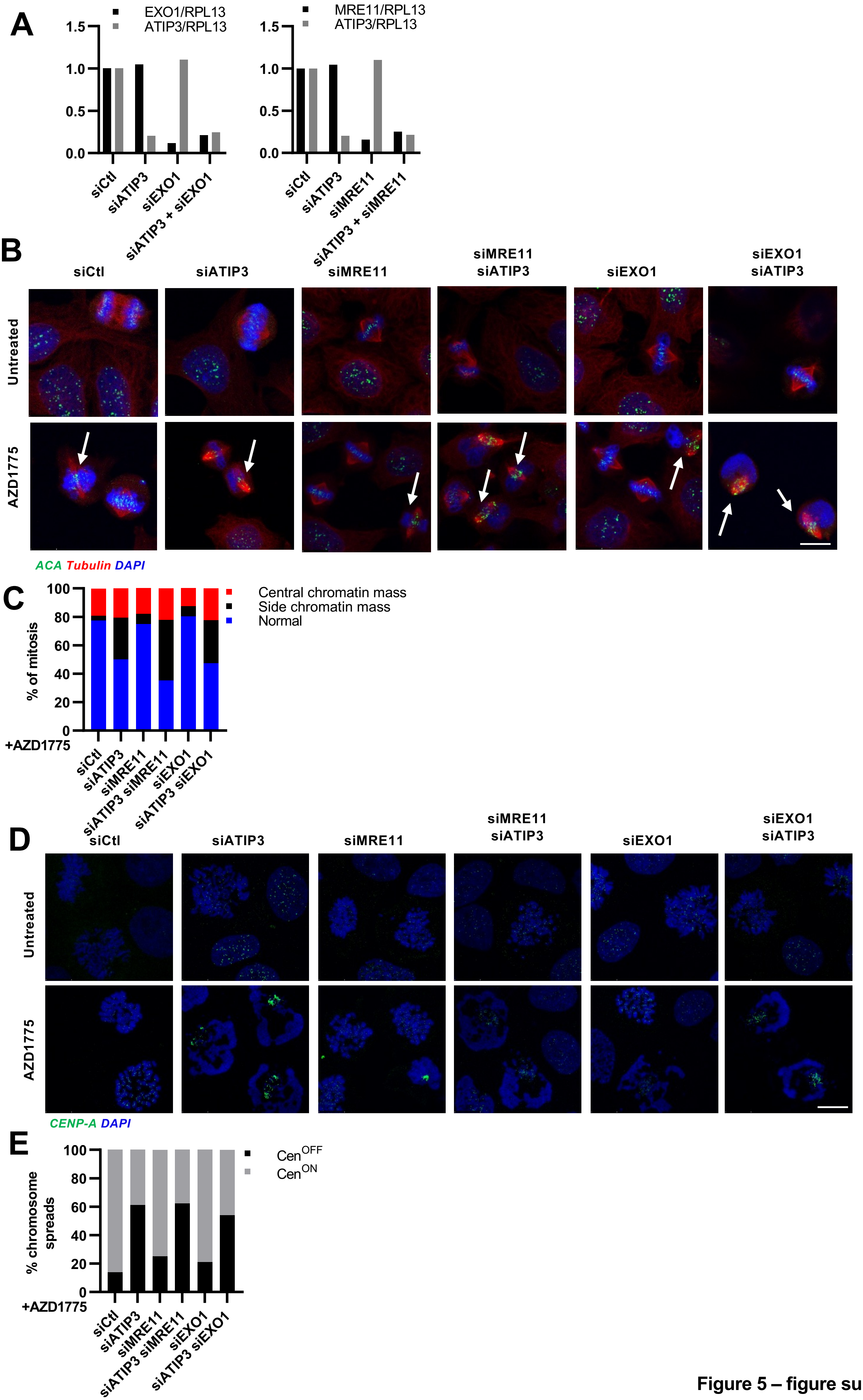


**Figure 5 – figure supplement 2**

### Supplemental Figure 5-3

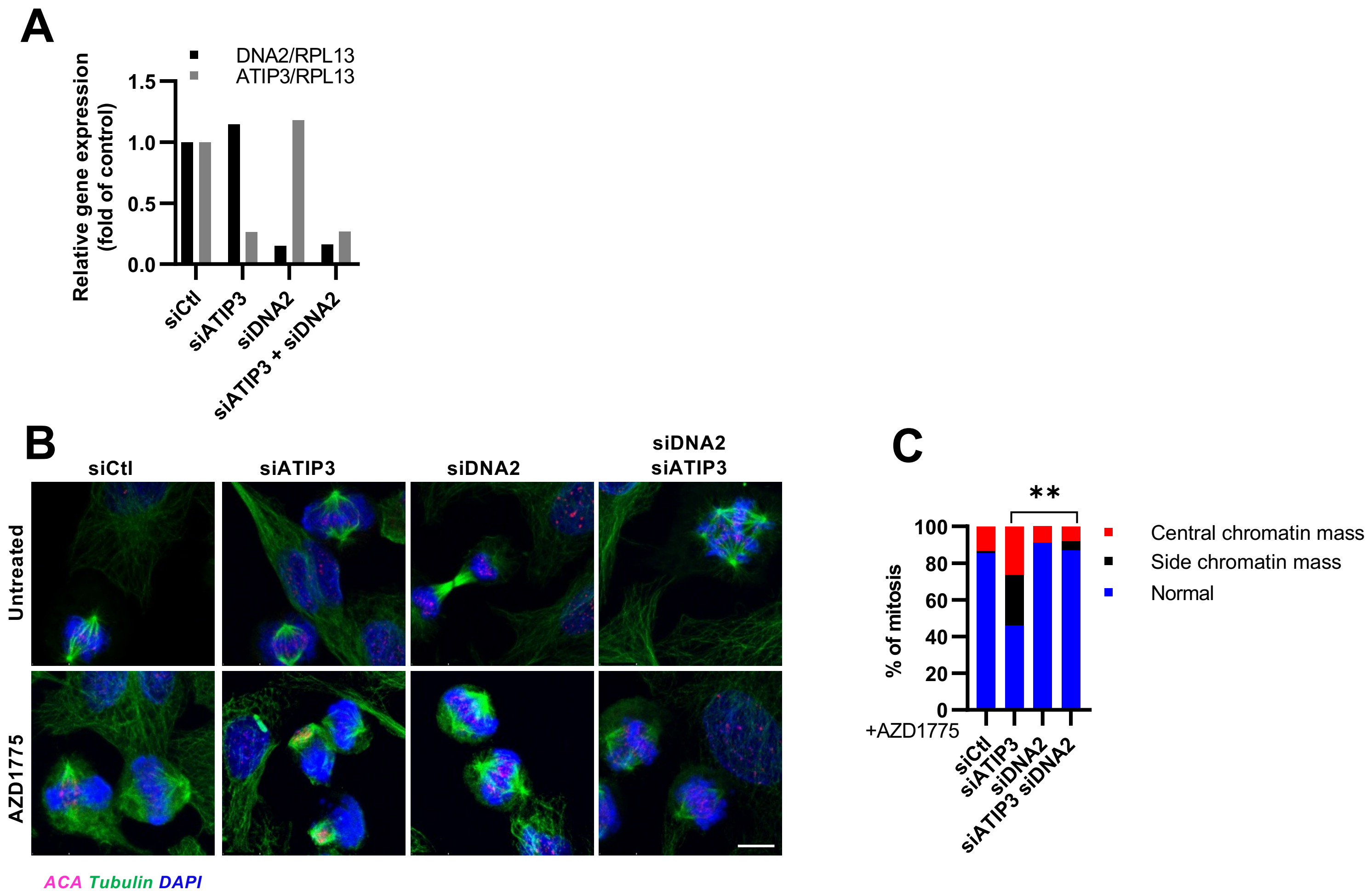


**Figure 5 – figure supplement 3**
